# Beyond Accuracy: Reliability-Aware Cross-Farm Evaluation of Dairy Cow Vocalization Models

**DOI:** 10.64898/2026.06.17.732832

**Authors:** Mayuri Kate, Suresh Raja Neethirajan

**Affiliations:** Faculty of Computer Science, Dalhousie University, Halifax, Canada; Faculty of Agriculture, Dalhousie University, Truro, Canada

**Keywords:** dairy cow vocalizations, self-supervised learning, acoustic unit discovery, proxy-state inference, cross-farm evaluation, probability calibration, selective prediction, precision livestock farming, animal welfare monitoring

## Abstract

Automated analysis of dairy cow vocalizations has largely relied on supervised classifiers evaluated within a single farm, a setting that inflates apparent performance and gives no measure of how far predictions can be trusted. We address this with a three-layer framework that separates acoustic structure discovery, proxy-state inference, and reliability assessment, evaluated on 569 annotated clips from three commercial dairy farms. A frozen self-supervised speech encoder, latent-space segmentation, and stability-guided clustering convert continuous recordings into discrete acoustic units without behavioral labels. Proxy-state signal is then tested under audio-only, audio-plus-context, and leave-one-farm-out (LOFO) protocols designed to separate transferable acoustic structure from farm-specific shortcuts. The results suggest that cross-farm generalizability differs substantially across biologically distinct vocalization categories. Non-vocal physiological sounds transfer across farms (LOFO macro-F1 = 0.763) and calibrate well (expected calibration error reduced from 0.087 to 0.023), whereas resource-related calls collapse to a majority-class baseline (macro-F1 = 0.500) and distress-related calls degrade under farm holdout. Selective prediction improves the retained-set score of the multiclass functional proxy (0.407 to 0.430), and an end-to-end convolutional baseline matches or exceeds the framework on raw accuracy for the easier targets yet yields a roughly two- to six-fold larger calibration error and offers no abstention. Random cross-validation consistently overstates cross-farm utility. These findings show that acoustic models for livestock monitoring require reliability-aware evaluation rather than flat classification.

## 1. Introduction

### 1.1 Dairy cow vocalization and the welfare-monitoring challenge

Dairy cattle are vocal animals whose calls carry information about physiological state, social motivation, reproductive readiness, and response to management events. This vocal expressiveness has long been recognized by animal scientists as a potential non-invasive window into welfare status: vocalizations can signal arousal, frustration, separation distress, and feeding-related motivation without requiring physical contact or restraint (Briefer, 2012; Manteuffel et al., 2004; Watts and Stookey, 2000). As precision livestock farming has expanded - integrating sensor networks, machine learning, and continuous monitoring into commercial dairy operations - acoustic monitoring has attracted growing interest as a practical route to automated welfare assessment (Neethirajan and Kemp, 2021; Neculai-Valeanu et al., 2025). Unlike contact-based sensors, fixed barn-mounted microphones are unobtrusive, low-cost, and capable of continuous data collection across entire herds without per-animal instrumentation. These properties make them technically attractive candidates for welfare monitoring at scale.

The scientific case for acoustic welfare indicators has been strengthened by a body of work linking vocal parameters to internal states. Fundamental frequency, call duration, spectral structure, and harmonic-to-noise ratio have all been associated with affective states ranging from feeding anticipation and estrus to separation distress and handling stress (Meen et al., 2015; Briefer, 2012). Scoping reviews confirm that bioacoustic monitoring now covers multiple welfare-relevant dimensions in farm animals, including pain, fear, frustration, and positive anticipatory states (Coutant et al., 2024). For dairy cattle specifically, research has linked vocalization rate to estrus climax, demonstrated individual vocal identity that persists across welfare-relevant contexts (Green et al., 2019), and shown that acoustic features differentiate behavioral categories under controlled conditions with accuracies exceeding 85% in several studies (Gavojdian et al., 2024; Kate and Neethirajan, 2026). Non-vocal physiological sounds like rumination, chewing, and respiration have similarly been shown to carry acoustically tractable information about feeding behavior and digestive health in grazing and housed cattle (Chelotti et al., 2020; Andriamandroso et al., 2016).

### 1.2 Limitations of current classification-based approaches

Despite this promise, automated dairy cow vocalization analysis has remained predominantly organized around supervised flat classification: assigning each audio clip or segment to a predefined category using models trained on labeled data from one or a small number of recording sites. Several structural limitations follow from this design. First, supervised classification systems learn representations that are entangled with the label set and the recording conditions used at training time. When deployed on new farms with different housing geometries, management routines, microphone placements, and acoustic environments, these models frequently show substantial performance degradation - a problem that has been identified as one of the central challenges in precision livestock farming AI (Kate and Neethirajan, 2026). Most published studies evaluate acoustic classifiers within a single cohort or under controlled laboratory conditions that do not reflect the acoustic heterogeneity of commercial barns (Gavojdian et al., 2024; Kate and Neethirajan, 2026), making it difficult to assess how much of the reported accuracy is generalizable and how much reflects farm-specific statistical regularities.

Second, existing approaches typically treat each clip as a fixed-length acoustic event and extract features from that clip as a whole, using either engineered descriptors (MFCCs, spectral centroid, fundamental frequency statistics) or end-to-end deep learning over raw waveforms. Neither approach explicitly asks whether the continuous vocal stream contains recurring structural units that could form a reusable symbolic vocabulary - one whose meaning might be conditioned by context rather than fixed. The notion that animal vocalizations, like speech, may contain sub-call structural elements that recur across behavioral contexts and whose interpretation depends on the surrounding sequence and situational setting has been suggested theoretically (Briefer, 2012) but has not been operationalized in dairy cow acoustic modeling. Without a stable, reusable structural vocabulary, every new classification task requires retraining from raw audio, and the learned representations cannot accumulate interpretable meaning across studies.

Third, the outputs of existing classifiers are almost invariably hard predicted labels - discrete class assignments without calibrated confidence estimates. A system that predicts "distress" with raw probability 0.51 and one that predicts it with raw probability 0.95 are reported identically in accuracy-based evaluations, yet they carry very different implications for downstream decision-making. The calibration literature has established that classification accuracy and probability reliability are distinct properties that can diverge substantially, particularly for imbalanced or context-sensitive targets (Guo et al., 2017; Silva Filho et al., 2023). The selective prediction literature has further shown that acknowledging uncertainty - withholding predictions when model confidence is low - can meaningfully improve the reliability of retained outputs in applied monitoring contexts (Geifman and El-Yaniv, 2017). Neither property has been systematically studied in dairy cow vocalization analysis.

These limitations are not three independent problems but three faces of the same structural gap. Existing livestock acoustic AI systems are organized around flat classification pipelines that are not stress-tested for cross-farm generalization and that provide no measure of prediction reliability. What is missing is a framework that separates acoustic representation, proxy-state inference, and trustworthiness as distinct scientific questions, and that evaluates each one under conditions where farm-specific shortcuts cannot rescue performance.

### 1.3 The present study

This study addresses that gap with a three-layer framework that moves dairy cow vocalization analysis beyond flat classification toward reliability-aware evidence modeling under real farm conditions. The novelty is not a new classifier, but a reproducible evaluation framework that separates three concerns prior work has conflated: how vocalizations are represented, how farm-specific context inflates apparent performance, and how far the resulting probabilities can be trusted. This distinction is critical because models that perform well under random cross-validation can fail under farm-held-out deployment conditions, which are the conditions a practical monitoring system must actually meet. Each layer is evaluated independently against its own question before its outputs pass to the next.

Figure 1 presents a high-level overview. Layer 1 converts continuous barn recordings into ordered sequences of discrete acoustic units using a frozen self-supervised speech encoder, latent-space segmentation, and stability-guided clustering, without any behavioral labels. Layer 2 tests whether these sequences carry proxy-state signal, contrasting audio-only, audio-plus-context, and leave-one-farm-out conditions to separate transferable acoustic structure from farm-specific shortcuts. Layer 3 then assesses how far the resulting predictions can be trusted, through target-specific calibration, selective prediction, and a model-behaviour dictionary. The layers are described in full in Sections 3–6.

**Figure 1.**
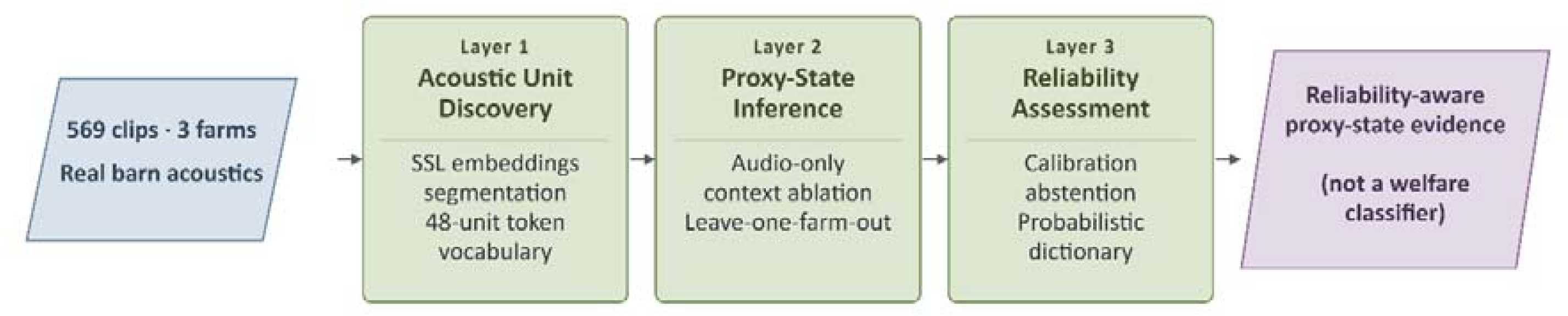
Overview of the three-layer analytical framework. Layer 1 converts real barn recordings into a reusable token vocabulary through self-supervised frame extraction, latent-space change-point segmentation, and unsupervised clustering. Layer 2 tests whether these token sequences carry proxy-state signal under audio-only, audio-plus-context, and leave-one-farm-out evaluation tracks. Layer 3 assesses prediction reliability through target-specific calibration, selective prediction, and model-behaviour dictionary construction.

The framework tests a single hypothesis: “Self-supervised acoustic unit discovery, combined with context-aware proxy-state modeling and reliability-aware interpretation, can move dairy cow vocalization analysis beyond flat classification toward cautious welfare-related evidence modeling”. The study does not produce a deployment-ready welfare classifier and does not claim access to biological ground truth. It produces a differentiated evidence profile - showing which proxy targets generalize across farms, under what conditions context adds interpretable information, and how reliably the framework’s probability estimates can be trusted.

### 1.4 Main contributions

Rather than introducing a new classifier, this study contributes a reproducible evaluation framework in which representation, context sensitivity, and probability reliability are treated as separate questions. The framework is designed for the condition that matters in practice: a model validated within one cohort may fail when an entire farm is held out. The contributions are methodological and evaluative:

1. Unsupervised acoustic unit discovery from real barn recordings. A frozen self-supervised encoder, latent-space change-point segmentation, and stability-guided clustering convert continuous barn audio into reproducible sequences of discrete acoustic units, without any behavioral labels. The frozen encoder is a deliberate choice that isolates framework-level behavior from task-specific tuning.
2. Context-audited cross-farm evaluation that exposes shortcut learning. A leave-one-farm-out protocol, contrasted with audio-only and audio-plus-context conditions, separates transferable acoustic structure from farm-specific correlations and identifies when a model relies on contextual shortcuts rather than generalizable signal. This addresses a gap in precision livestock AI, where within-cohort validation predominates and cross-farm robustness is rarely tested.
3. Reliability-aware prediction through calibration and selective prediction. Target-specific post-hoc calibration and abstention separate discriminative performance from probability quality, showing when model confidence is trustworthy and when a prediction should be deferred. While both techniques are established, their systematic, target-by-target application to livestock acoustic modeling is, to our knowledge, new to this domain.
4. Evidence-based interpretation of which proxy targets generalize and which collapse. Cross-farm evaluation on real commercial data shows that generalizability differs substantially across biologically distinct vocalization categories: non-vocal physiological sounds transfer across farms, whereas resource- and distress-related proxies degrade or collapse under farm holdout. This demonstrates that performance gains observed under standard within-cohort validation may not reflect true cross-farm utility.

## 2. Materials, Dataset, and Study Framing

### 2.1 Dataset origin and collection

The corpus used in this study was collected and described in full by Kate and Neethirajan (2026) and is briefly summarized here as it provides the acoustic foundation for all analyses. The dataset comprises 569 curated audio clips spanning nine main behavioral categories and 39 subcategories, recorded across three commercial dairy farms in Sussex County, New Brunswick, Canada. Data collection took place over three consecutive days (5–7 May 2025), with one farm visited per day between 09:00 and 18:00. The farms are referred to throughout as Farm 1, Farm 2, and Farm 3. Farm 1 housed 57 Holstein cows and contributed 174 clips. Farm 2, a mixed Holstein-Jersey herd of 207 animals, contributed 265 clips. Farm 3 housed 160 Holstein cows and contributed 120 clips, the full cross-tabulation of clip counts by farm and behavioral category is given in Supplementary Table S2. In total, 65 raw audio files representing approximately 90 hours of recordings (65.6 GB) were collected across the three sites. Table 1 summarises the audio dataset.

**Table 1.** Dataset summary: recording sites, clip counts, barn zones, and recording setup.

|  | **Farm 1** | **Farm 2** | **Farm 3** | **Total** |
| --- | --- | --- | --- | --- |
| Herd size | 57 Holstein | 207 Holstein & Jersey | 160 Holstein | 424 cows |
| Clips contributed | 174 | 265 | 120 | 559 (farm-identified) |
| Recording date | 5 May 2025 | 6 May 2025 | 7 May 2025 | 3 consecutive days |
| Barn zones | Feeding, Drinking, Milking, Resting | Feeding, Drinking, Milking, Resting | Feeding, Drinking, Milking, Resting | 4 zones total |
| Raw files | 21 | 21 | 23 | 65 |
| Raw recording duration | ~30 h | ~29 h | ~31 h | ~90 h |
| Raw data size | ~21.5 GB | ~22.0 GB | ~22.1 GB | 65.6 GB |

*Note: Ten additional clips in the corpus lack confirmed farm identification and are excluded from the farm-identified total of 559. All 569 clips were used in within-cohort analyses; only the 559 farm-identified clips were used in leave-one-farm-out evaluation.*

### 2.2 Recording design and equipment

Recordings were made using a multi-microphone array deployed at fixed positions within each barn zone, allowing simultaneous capture from multiple spatial locations within the same farm (Kate and Neethirajan, 2026). The setup combined three complementary equipment configurations. Directional shotgun microphones (Sennheiser MKH 416) paired with multi-channel portable recorders (Zoom F6) were deployed in feeding alleys and resting areas to capture high-fidelity close-range vocalizations while suppressing off-axis barn noise. Supercardioid shotgun microphones (RØDE NTG-2) paired with handheld recorders (Zoom H4n Pro) were used at drinking troughs and milking parlors for flexible mobile coverage. A Wildlife Bioacoustics autonomous logger was deployed in resting areas for unattended continuous monitoring. GoPro action cameras were mounted above each barn zone, providing synchronized video for behavioral context cross-referencing. Microphones were positioned approximately one meter above cow head height and oriented toward zone centers, placing them within approximately 0.5–3 m of vocalizing animals as cows moved naturally through the monitored areas (Kate and Neethirajan, 2026).

Across the 569 curated clips, three main barn zones dominated the corpus: Feeding (268 clips, 47.1%), Resting (144 clips, 25.3%), and Drinking (62 clips, 10.9%). The Milking zone contributed 54 clips (9.5%) and the Water Station zone 31 clips (5.5%); the distribution of clip counts across barn zones and behavioral categories is detailed in Supplementary Table S3. Clip durations followed a right-tailed distribution with a mean of 21.04 seconds, a median of 13.81 seconds, and a range of 2.80 to 444.9 seconds (10th–90th percentile: 11.58–30.77 seconds, standard deviation 28.68 seconds). The extended tail reflects the dataset’s behavioral diversity - sustained rumination sessions, prolonged estrus episodes, and extended social exchanges naturally produce much longer clips than brief single calls - and is a structural feature of an ecologically realistic corpus rather than a data quality concern (Kate and Neethirajan, 2026).

### 2.3 Behavioral annotation scheme

Raw recordings were preprocessed using spectral noise profiling, band-pass filtering (50–1,800 Hz), and adaptive spectral denoising to suppress barn machinery noise while preserving the bovine vocal band (Kate and Neethirajan, 2026). Clips were manually segmented using Raven Lite 2.0, with onset and offset boundaries verified acoustically in Praat against pitch contour and formant continuity criteria. Each clip was then independently annotated by two trained observers, who assigned a main behavioral category, a subcategory, an emotional context label, and a confidence score, with access to co-recorded video where needed to resolve ambiguity.

Inter-annotator agreement on main-category labels across a stratified subset of 150 clips yielded a Cohen’s κ of 0.884 with 90.0% raw agreement. For the more fine-grained subcategory labels, κ was 0.699 with 71.3% raw agreement. Final consensus labels were derived through joint review by both annotators and the lead investigator, with agreement between consensus and individual annotator labels reaching κ = 0.930 and κ = 0.954 for main categories, indicating an auditor-validated annotation set (Kate and Neethirajan, 2026). The annotation scheme was informed by ethological principles, linking call types to proximate physiological and environmental triggers and to their broader communicative functions in line with contemporary animal welfare frameworks.

The nine main behavioral categories in the original annotation scheme, and their clip counts in this corpus, are shown in Supplementary Table S1. Subcategory coverage spans 39 distinct subcategories across the 569 clips, with a strongly right-skewed class distribution. The class imbalance in the dataset, which mirrors the natural frequency of these behaviors on commercial farms, has direct implications for the proxy-state recoding described in Section 2.5.

### 2.4 Available metadata and its constraints

Each clip is accompanied by the following structured metadata fields: farm identity, barn zone, microphone model, microphone placement context, recording date, recording day, main behavioral category, and subcategory. These fields provide the contextual backbone for the context-aware inference experiments in Layer 2.

Several important metadata fields are absent from this corpus, and their absence directly shapes what can and cannot be claimed in the analyses that follow. No synchronized physiological measurement is available - cortisol levels, heart rate, or other direct welfare indicators were not recorded alongside the vocalizations. No individual cow identification is present - clips cannot be attributed to specific animals, and repeated vocalizations by the same individual cannot be distinguished from vocalizations by different animals. No feeding or milking schedules were recorded in a form that could be precisely aligned with clip timestamps. No activity sensor data, body condition scores, or parity information accompany the recordings. Video was collected in parallel but was used only for annotation cross-referencing during curation; it was not processed into quantitative behavioral variables available for modeling.

These absences are not exceptional for a real-barn vocalization corpus. As Kate and Neethirajan (2026) note, the practical difficulty of obtaining temporally aligned physiological measurements in commercial farm settings is a widely acknowledged limitation across the bovine bioacoustics literature. The dataset’s strength lies precisely in its ecological realism - authentic barn acoustics, multiple farms, diverse behavioral contexts - which comes at the cost of the controlled synchrony achievable only in laboratory settings. The analytical framework in this study was designed from the outset to work honestly within these constraints rather than around them.

### 2.5 Why proxy-state framing is necessary

The behavioral annotation categories in the corpus record what was observable and audible at the time of recording. They do not constitute validated welfare ground truth. A clip labeled Distress & Pain represents a human annotator’s judgment that the acoustic event and its behavioral context were consistent with a distress-related call; it does not independently verify the animal’s physiological state, arousal level, or welfare score at that moment. Similarly, an Estrus & Mating Behavior label indicates that the acoustic and behavioral profile was consistent with estrus-associated calling, not that an estrus event was independently confirmed through hormonal assay. The same acoustic structure - a moderately loud call recorded in the milking parlour - could be consistent with pre-milking anticipation, handling stress, or maternal separation depending on context not fully captured in the metadata.

This gap between behavioral annotation and validated welfare state is the scientific motivation for proxy-state framing throughout this study. Rather than treating the annotation categories as biological ground truth and building models to recover them, the framework treats them as operational proxies - functionally defined groupings that allow hypotheses about acoustic signal to be tested without asserting that the label encodes direct welfare knowledge. This is consistent with the broader shift in animal welfare science from simple behavioral indicators toward probability-based, context-conditioned evidence profiles (Briefer, 2012; Coutant et al., 2024).

For the inference experiments in Layer 2, the nine original annotation categories and 39 subcategories were consolidated into four proxy-state targets through a biologically motivated recoding that preserved interpretable functional groupings while reducing the long-tail sparsity of the full subcategory space. The recoding logic, class compositions, and clip counts are given in Section 5 and Table 3.

**Table 2.** Summary of the three-layer framework: scientific role, methodology, and outputs.

| Layer | Scientific role | What it receives | Core methodology | Principal outputs |
| --- | --- | --- | --- | --- |
| Layer 1:<br>Acoustic<br>Foundation | Discover stable acoustic unit structure without supervision | 569 audio clips and structural recording metadata | Frozen SSL frame extraction, latent-space change-point segmentation with refinement sweep, stability-based K-study clustering (K=48), token assignment, context bias audit | 48-unit acoustic vocabulary, 4,764 token events, ordered clip-level sequences, bias-audited unit inventory |
| Layer 2:<br>Proxy-State<br>Inference | Test whether discovered structure carries proxy-state signal and whether context conditions its interpretation | Token sequences and recording-context metadata from Layer 1 | Composition and sequence-transition feature engineering, audio-only / audio+context / leave-one-farm-out evaluation, full and high-bias-filtered vocabulary variants | Per-target probability estimates, discriminative benchmarks, context-gain diagnostics, cross-farm spread analysis |
| Layer 3:<br>Reliability<br>Assessment | Establish how trustworthy the proxy-state | Layer 2 prediction outputs | Target-specific post-hoc calibration, selective prediction and | Calibrated probability estimates, abstention maps, reliability- |
|  | probabilities are and under what conditions they can be interpreted | across fixed benchmark bundles | abstention analysis, model-behaviour dictionary construction, confidence-stratified case review | tiered token dictionary, confidence-stratified case catalog |

**Table 3.** Proxy-state target definitions, class compositions, and interpretive roles.

| Proxy state | Original categories merged | Clips (n) | Binary target | Interpretive role in this study |
| --- | --- | --- | --- | --- |
| physiological_nonvocal | Non-Vocal Sounds | 142 | is_nonvocal | Flagship target: acoustically most distinct and farm-generalizable |
| ingestion_resource | Feeding & Hunger Related; Water & Thirst Related | 169 | is_resource_related | Stress-test target: resource-access behavior is likely farm-routine dependent |
| reproductive_estrus | Estrus & Mating Behavior | 127 | - | Multiclass component; retained within proxy_functional_v1 |
| stress_discomfort | Distress & Pain; Milking & Handling; Environmental & Situational | 67 | is_distress_related | Biologically meaningful but context-sensitive; mixed cross-farm behavior |
| social_contact | Social Recognition & Interaction; Maternal & Calf Communication | 64 | - | Multiclass component; acoustically overlapping with resource and stress categories |

This proxy-state framing is methodologically conservative by design. It avoids asserting that the model recovers a ground-truth welfare state and instead asks whether acoustic token sequences carry statistically reliable signal that discriminates among these functional proxies - under audio-only conditions, under context-conditioned conditions, and under the more demanding cross-farm evaluation that tests whether the signal generalizes beyond the farms present in the training data.

## 3. Overall Framework

### 3.1 Framework rationale

The study is organized as a layered analytical framework rather than a single end-to-end classifier, for two reasons. The first is biological: dairy cow vocalizations are context-dependent signals whose meaning cannot be read from acoustic form alone, so a system that treats them as fixed-label stimuli risks predictions that are accurate within a recording cohort but uninformative outside it (Briefer, 2012; Coutant et al., 2024). The second is epistemological: without synchronized physiological or behavioral ground truth, the defensible objective is not welfare-state prediction but a measured account of how far acoustic evidence alone can carry cautious proxy-state inference, under what conditions context helps, and how reliably the resulting predictions can be interpreted.

These constraints motivate three layers that address progressively higher-order questions (Figure 2). The first asks whether vocalizations contain recurring acoustic structure that can be represented stably and without supervision. The second asks whether that structure carries proxy-state meaning, and whether context conditions it in ways that survive cross-farm evaluation. The third asks whether the resulting probabilities are trustworthy enough to support evidence statements. The layers are not detached experiments; each constrains the claims the next can make.

**Figure 2.**
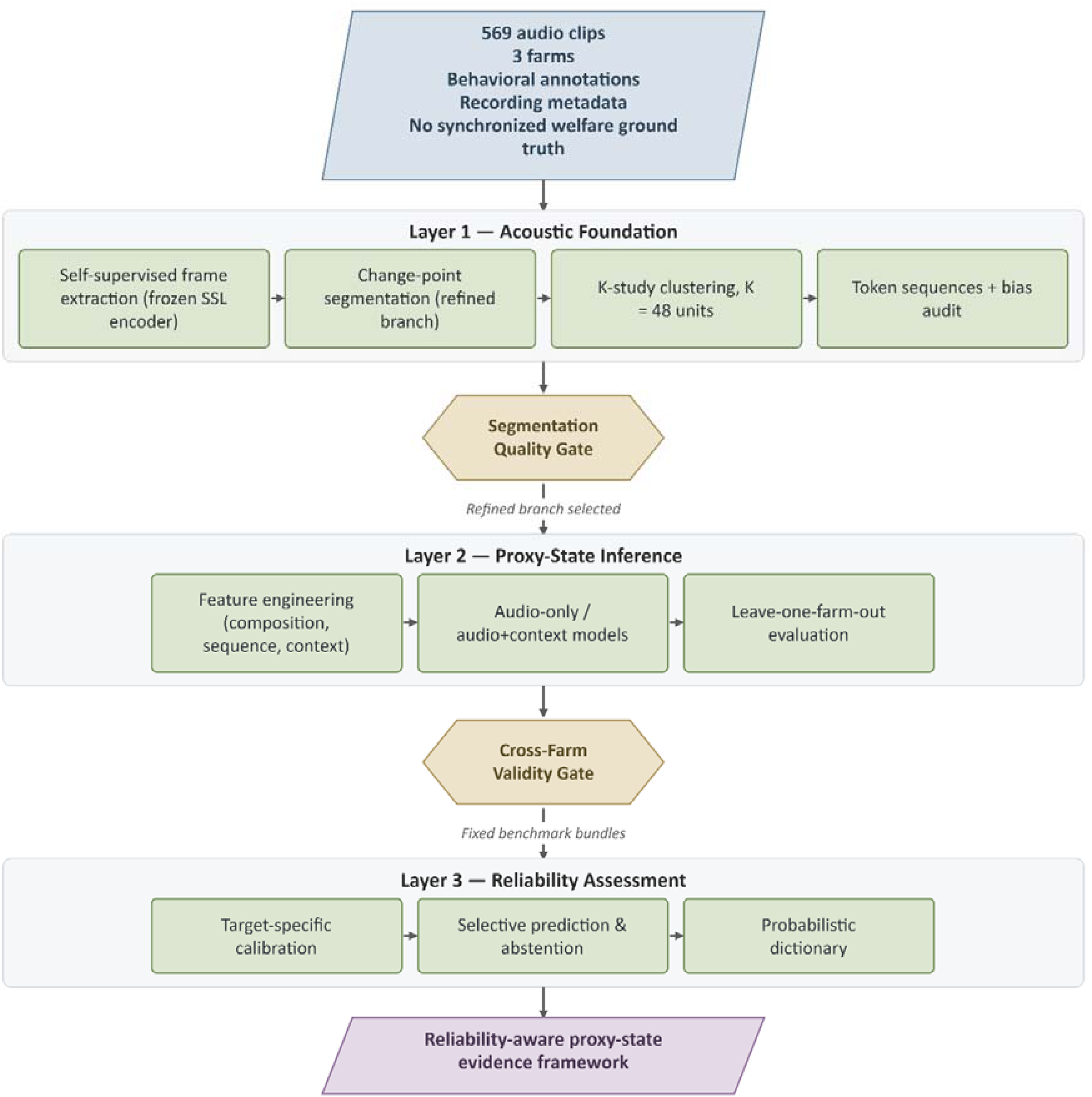
Architecture of the three-layer framework showing the forward data flow and the parallel evaluative flow. Each layer receives fixed inputs from its predecessor and produces auditable intermediate outputs. The three evaluation tracks - audio-only, audio-plus-context, and leave-one-farm-out - are maintained as parallel benchmarks from Layer 2 onward, enabling differentiated assessment of each proxy target’s robustness.

### 3.2 Architectural logic and methodological traceability

Two design decisions support this cumulative reading. First, the framework maintains a forward data flow - acoustic unit discovery, token-sequence construction, proxy-state prediction, and reliability assessment - alongside a parallel set of methodological gates (segmentation adequacy, inventory stability, context-gain, cross-farm robustness, calibration benefit, and abstention utility) that keep each performance claim tied to an explicit check rather than letting downstream accuracy conceal upstream weakness. Second, three evaluation tracks are preserved throughout: an audio-only reference establishes what acoustic structure alone recovers, an audio-plus-context reference tests whether meaning is context-conditioned, and a leave-one-farm-out benchmark tests whether those gains survive once farm-specific shortcuts are removed. All intermediate outputs - segmentation boundaries, cluster assignments, token sequences, dictionary scaffolds, and benchmark bundles - are fixed as auditable artefacts at each transition, so every layer inherits a stable input rather than a moving target.

This layered design also supports a differentiated reading of the results: targets that generalize across farms, calibrate well, and benefit from abstention emerge as robust, while targets that collapse under farm holdout remain visible as boundary cases rather than being absorbed into a single pooled score. In a domain where variation in recording conditions and biological context is unavoidable, that differentiated evidence profile is more informative than one aggregate accuracy figure.

## 4. Layer 1: Acoustic Foundation

### 4.1 Overview and design rationale

Layer 1 converts raw audio into ordered sequences of discrete acoustic tokens whose structure is discovered without supervision, predefined labels, or any downstream classification signal. The aim is not to classify sounds but to build a stable, reusable acoustic vocabulary - units that recur across clips, whose ordering within each clip forms the input to Layer 2. This departs from fixed-window, feature-engineered tokenization, which imposes artificial boundaries on continuous acoustic change and produces tokens reflecting windowing choices rather than the signal itself.

The pipeline proceeds through six stages - frame-level SSL embedding extraction, baseline segmentation, segmentation refinement and candidate selection, clustering-based unit discovery, token assignment, and context bias auditing - each producing preserved, traceable outputs (Figure 3).

**Figure 3.**
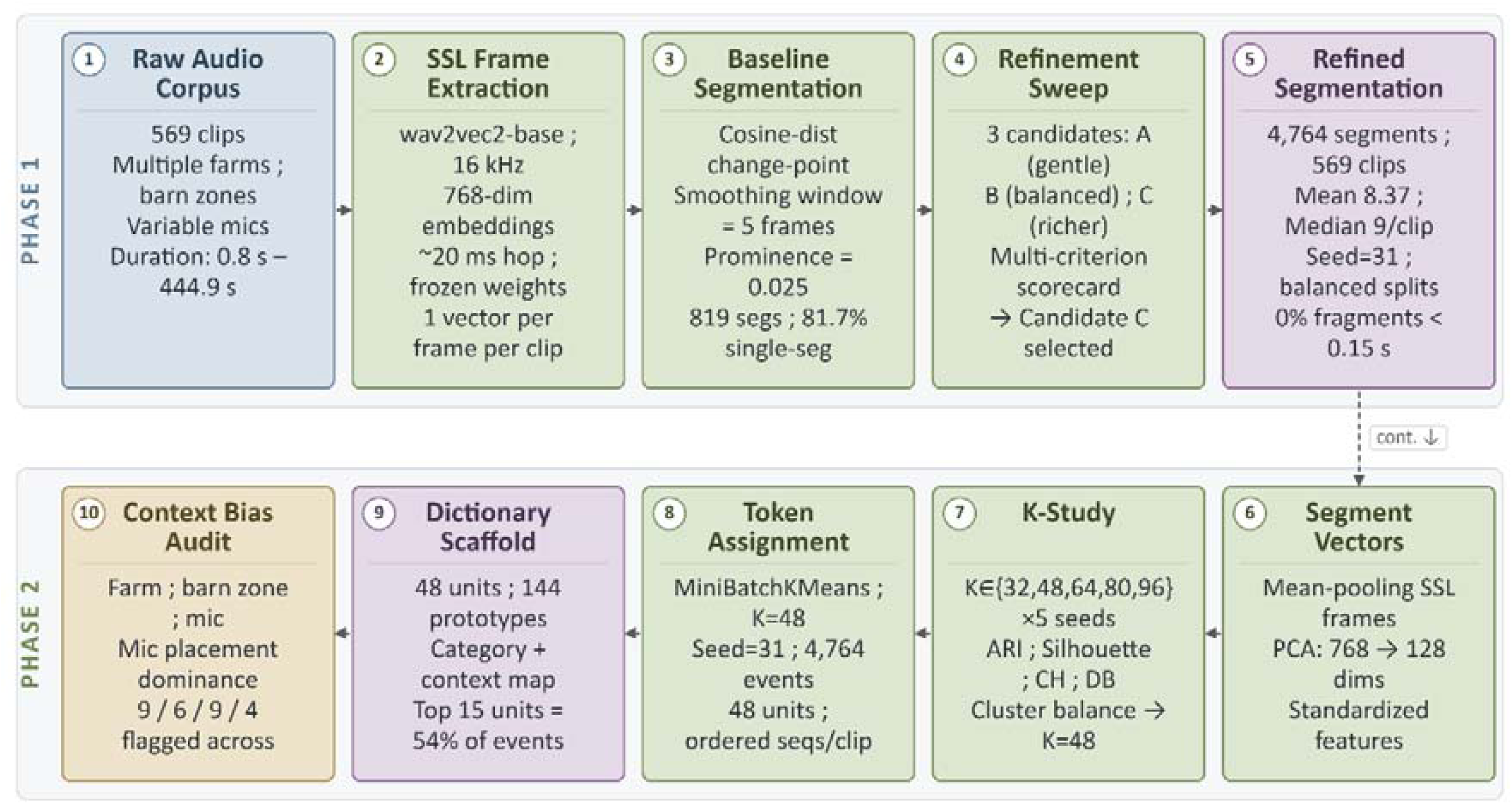
Layer 1 pipeline from raw audio to bias-audited token vocabulary. Processing stages proceed left to right: SSL frame extraction using frozen wav2vec 2.0, latent-space change-point segmentation with the refined Candidate C configuration, PCA-reduced segment vector construction, stability-guided K-study clustering at K = 48, ordered token sequence assignment, and context bias auditing. The output is a 48-unit acoustic vocabulary and 4,764 token events across 569 clips.

### 4.2 Dataset and audio preparation

Layer 1 took the 569 clips from three commercial farms (Section 2). Their durations are highly heterogeneous (Figure 4; per-category duration statistics in Supplementary Table S4) - extended rumination, brief single calls, long milking-zone recordings - which motivated frame-level extraction rather than clip-level pooling: a single pooled vector per clip would discard all within-clip temporal structure and make segmentation impossible.

**Figure 4.**
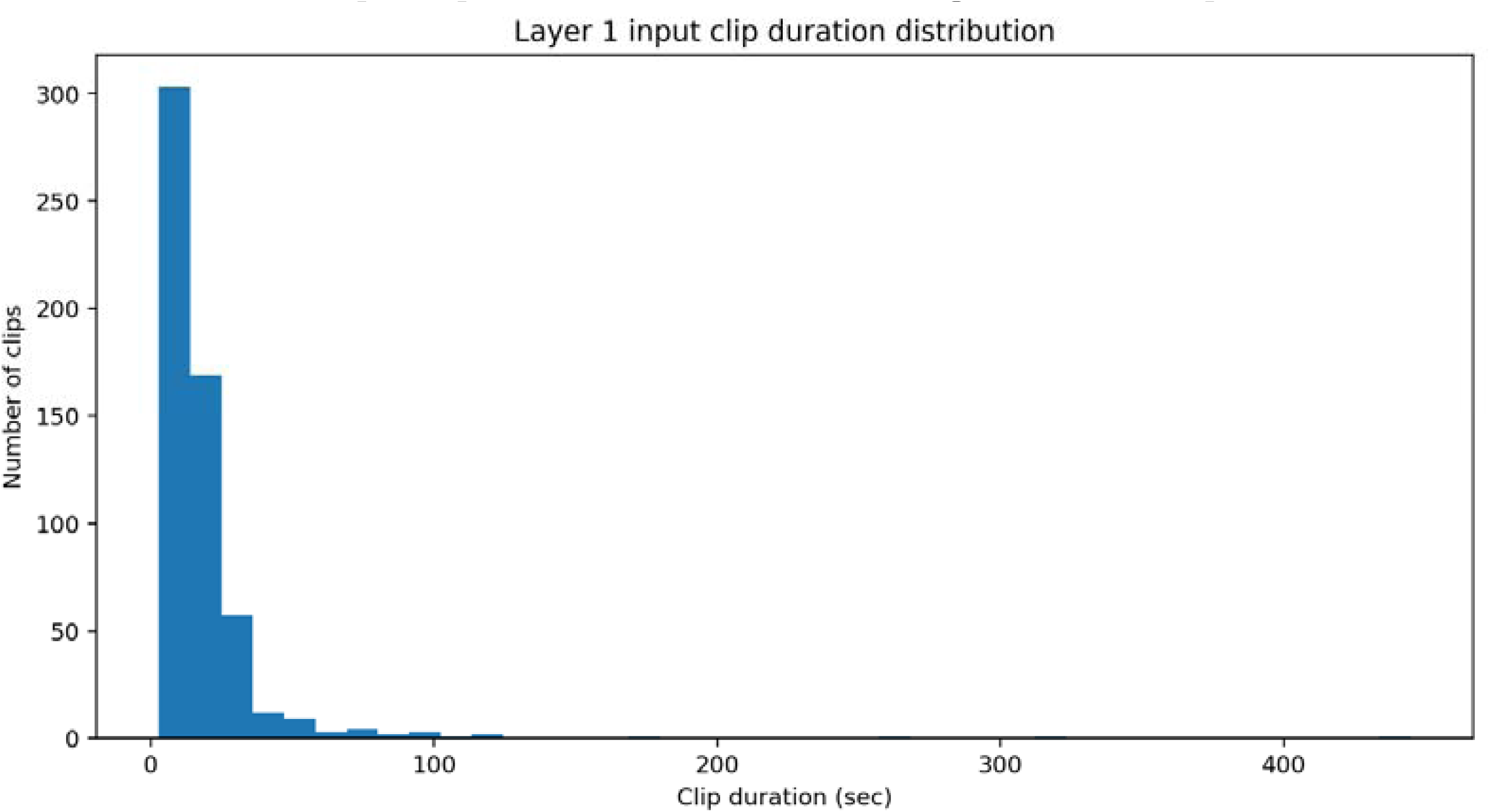
Distribution of clip durations across the 569 curated audio clips. The right-tailed distribution reflects the behavioral diversity of the corpus, ranging from brief single calls (minimum 2.80 s) to sustained rumination and estrus episodes (maximum 444.9 s). The median duration is 13.81 s, with a mean of 21.04 s. This duration heterogeneity motivated frame-level feature extraction rather than clip-level pooling.

All audio was loaded as mono waveform and resampled to 16,000 Hz. Peak normalization was applied prior to SSL feature extraction to reduce the influence of recording-level gain differences across farms and microphone models.

### 4.3 SSL frame-level representation

Frame-level embeddings were extracted with the frozen wav2vec2-base model (Baevski et al., 2020), a self-supervised encoder pretrained on 960 hours of unlabeled English speech (LibriSpeech), comprising a convolutional feature encoder (≈20 ms per frame) and a Transformer context encoder. No weights were updated and no fine-tuning was performed. This frozen-encoder choice is deliberate: it isolates the contribution of the downstream framework from task-specific tuning and utilizes general-purpose acoustic representations as a structural foundation, rather than training a domain-specific encoder that would require labelled cow data and would not generalize beyond it.

Each clip yielded 768-dimensional frame embeddings (mean ≈1,051 frames, median ≈691) at a ≈20 ms hop, the resolution needed for change-point detection. Speech-pretrained SSL models transfer effectively to bioacoustics even when the target species differs substantially from the pretraining domain (Hagiwara, 2023; Sarkar and Magimai-Doss, 2025). The objective of this study was to evaluate the analytical framework and its cross-farm reliability, rather than to maximize the quality of the underlying acoustic representation. A stable, extensively benchmarked backbone was therefore selected intentionally, so that observed performance reflects the behaviour of the framework rather than encoder-specific tuning. Animal-specific or bioacoustic self-supervised encoders may yield richer species-tuned representations, but evaluating them is an orthogonal question to the framework-level assessment pursued here. A speech backbone is not specialised for bovine acoustics, but it provides a reproducible representation that does not depend on scarce labelled cow data; substituting an animal-specific encoder such as AVES (Hagiwara, 2023) is a deliberate future direction, not a requirement for the present framework-level evaluation (Section 8).

### 4.4 Latent-space change-point segmentation

#### 4.4.1 Baseline segmentation and its limitations

Segmentation detected temporal boundaries in the SSL frame sequence using a latent-space change-point method: instead of fixed windows, it locates points where local acoustic content changes abruptly, using cosine distance between consecutive frame embeddings as the change signal. The procedure (Figure 5) smooths frame embeddings (uniform filter, 5 frames), computes and smooths frame-to-frame cosine distances (3 frames), detects boundaries as peaks via a prominence threshold and a MAD-based height filter robust to clip-level change-score scale, and merges sub-threshold fragments into adjacent segments. The baseline configuration (prominence 0.025, MAD multiplier 3.0, minimum inter-peak distance 0.20 s, minimum segment 0.25 s) produced only 819 segments, with 465 of 569 clips (81.72%) assigned a single segment - degenerate for token-sequence modeling - prompting a systematic refinement.

**Figure 5.**
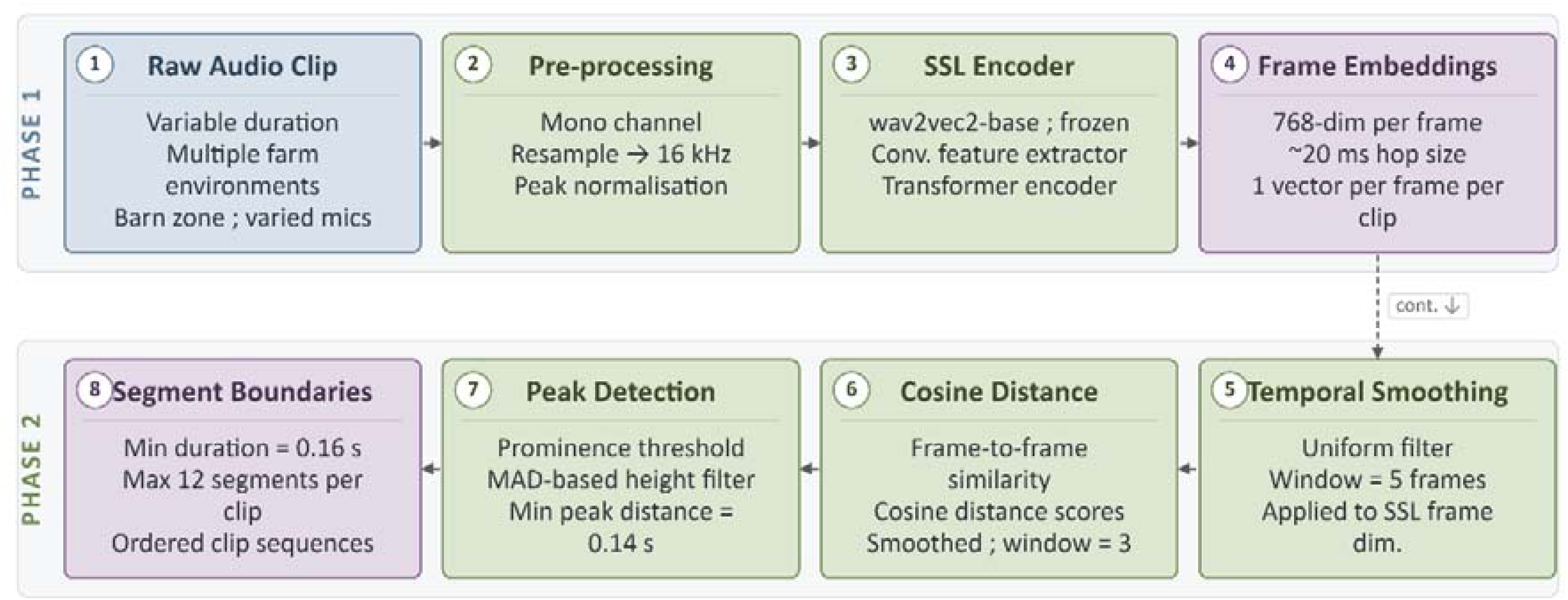
Two-phase processing flow from raw audio to segment boundaries under the refined Candidate C configuration. Phase 1 (top, left to right): raw audio clips are pre-processed to mono channel with 16 kHz resampling and peak normalization, passed through the frozen wav2vec2-base encoder (convolutional feature extractor followed by Transformer encoder), and converted to 768-dimensional frame embeddings at approximately 20 ms hop size. Phase 2 (bottom, right to left): frame embeddings are temporally smoothed (uniform filter, window = 5 frames), frame-to-frame cosine distances are computed and smoothed (window = 3), peaks are detected using prominence thresholds and a MAD-based height filter (minimum peak distance = 0.14 s), and segment boundaries are finalized with a minimum duration of 0.16 s and a maximum of 12 segments per clip.

#### 4.4.2 Segmentation refinement sweep

A controlled sweep of three alternative parameter configurations was conducted. All three candidates used the same general algorithm but applied progressively lower detection thresholds to recover more within-clip boundaries. The configurations, labeled Candidate A (gentle split), Candidate B (balanced split), and Candidate C (richer sequences), are summarized in Supplementary Table S8.

Candidate selection used a five-criterion weighted scorecard combining single-segment improvement, fragment control, downstream clustering quality, sequence richness, and confound control, with penalty terms for fragmentation and increased context locking. The full criteria, weights, and penalties are given in Supplementary Methods S1.

Candidate C scored highest (0.600, versus 0.351 and 0.100 for Candidates B and A) and was adopted as the official refined segmentation. It produced 4,764 segments across the 569 clips, a mean of 8.37 segments per clip, only 6 single-segment clips (1.05%), and no fragments below 0.15 s. All downstream Layer 1–3 analyses used this segmentation.

The refined segmentation produces sequences of acoustic segments within each clip that reflect within-clip structural change in the SSL embedding space, rather than arbitrary windowing. Each clip now yields between 1 and 12 segments, with most clips contributing 6–12 segments (Figure 6). This richness of within-clip structure is what enables token-sequence modeling in Layer 2.

**Figure 6.**
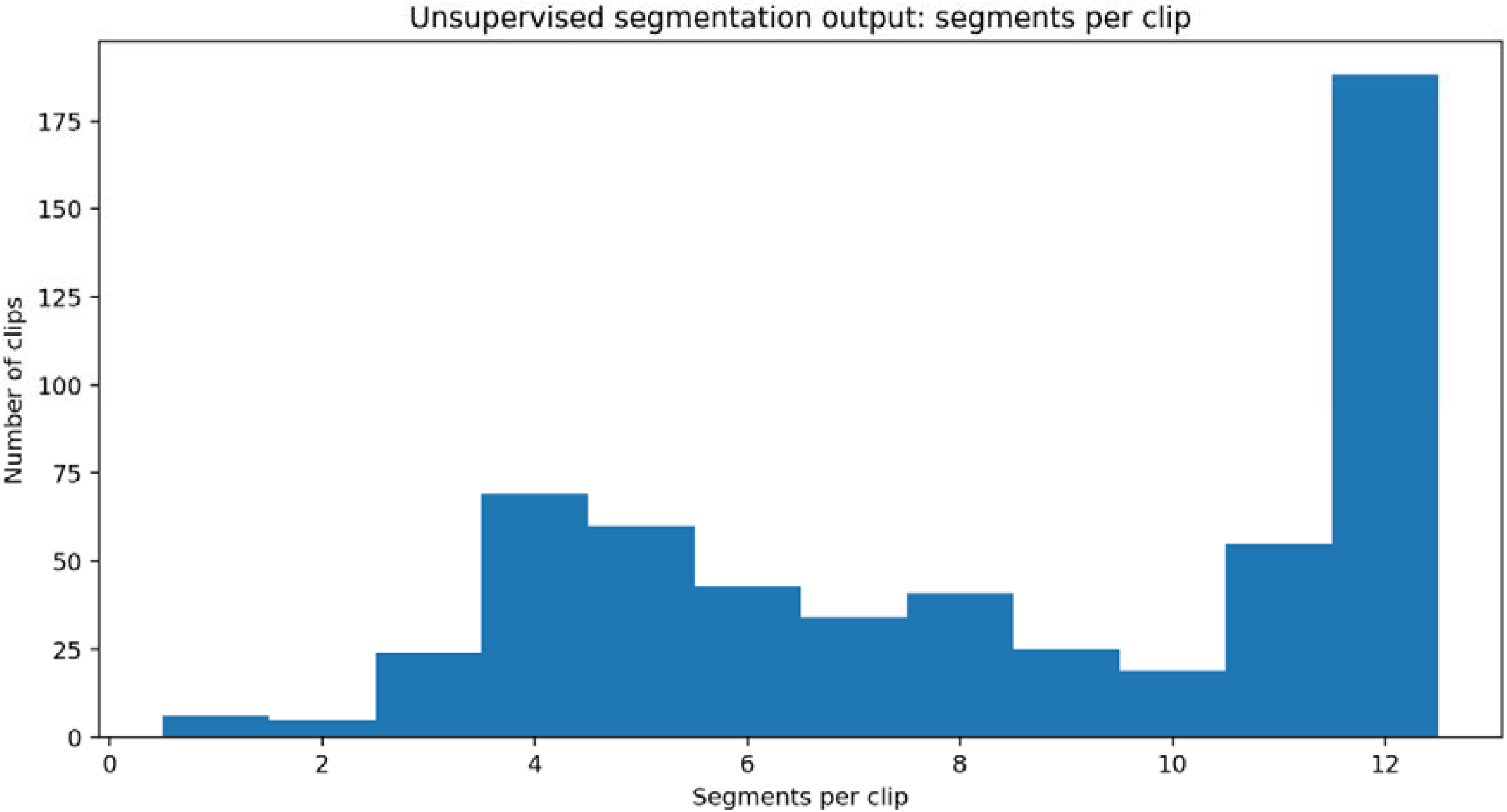
Distribution of segments per clip under the refined Candidate C segmentation. The majority of clips contain between 6 and 12 segments (median = 9), with only 6 clips (1.05%) retaining a single segment. The maximum of 12 segments per clip is enforced by the configuration ceiling. This within-clip structural richness enables token-sequence modeling in Layer 2.

### 4.5 Segment-level vector construction

Each segment was represented by mean-pooling the SSL frame embeddings within its boundaries (more robust than first- or last-frame selection). The 768-dimensional vectors were standardized (zero-mean, unit-variance) across all 4,764 segments, then reduced to 128 components by PCA - removing low-variance dimensions and lowering K-study cost - and used as clustering input.

### 4.6 Acoustic unit inventory selection: K-study

The vocabulary size K was selected empirically rather than assumed. Five values (K ∈ {32, 48, 64, 80, 96}) were each evaluated over five MiniBatchKMeans runs with different random seeds, and reproducibility was quantified by the mean pairwise Adjusted Rand Index (ARI) across runs. Internal cluster-quality and balance metrics were also computed and combined into a composite score that penalised singleton or very small clusters. Full settings and per-K metrics are reported in Supplementary Methods S2 and Supplementary Table S9, with the stability profile in Figure 7.

**Figure 7.**
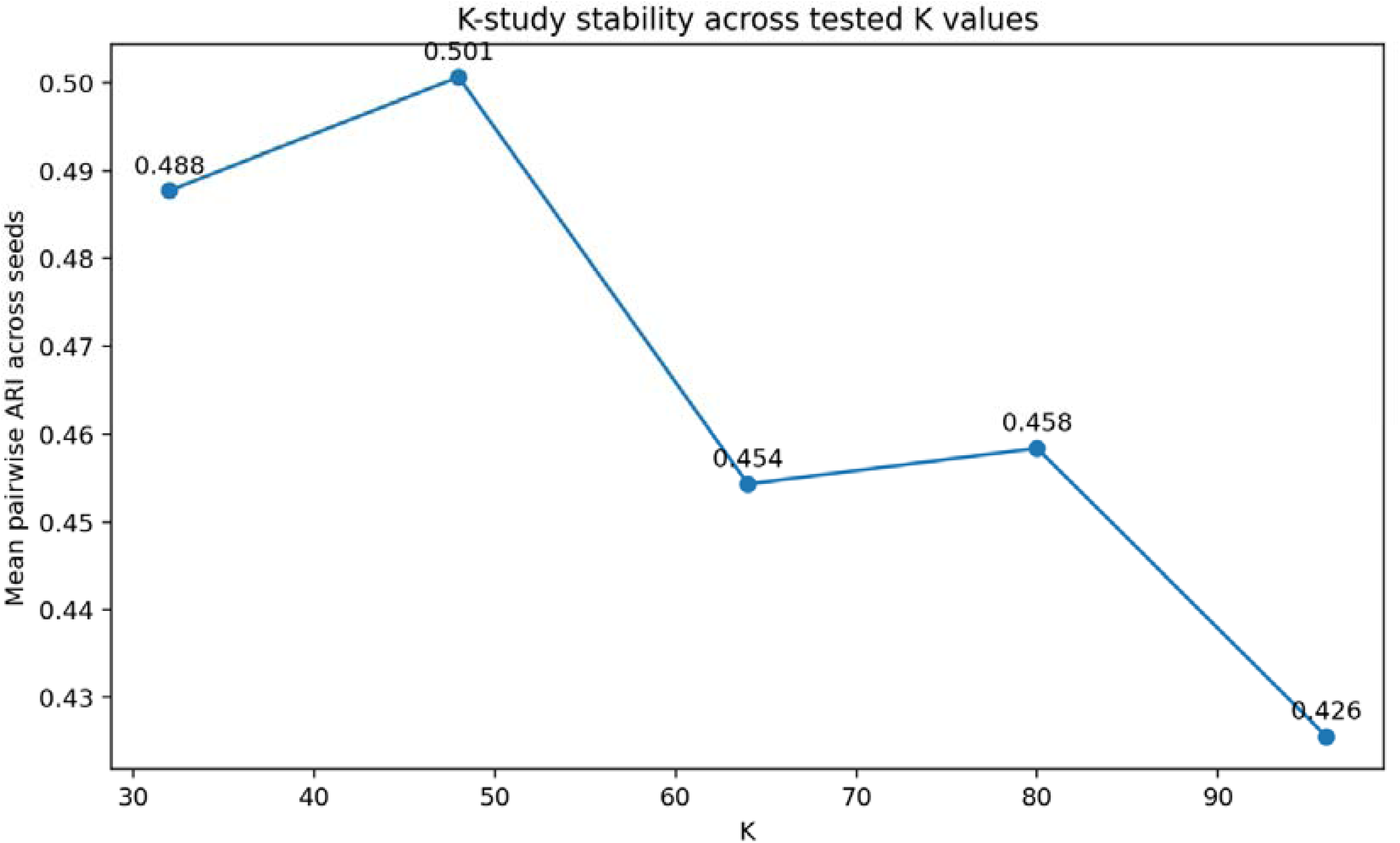
Pairwise Adjusted Rand Index (ARI) stability across five clustering seeds for each candidate K value. K = 48 achieved a mean pairwise ARI of 0.501 with moderate seed-to-seed variation. Higher K values (64, 80, 96) showed progressively lower ARI and increased instability, while K = 32 offered higher stability at the cost of reduced vocabulary resolution.

K = 48 was selected as the unit inventory. It gave a mean pairwise ARI of 0.501 - moderate but acceptable reproducibility for an unsupervised problem of this complexity - with zero singleton clusters and a normalized cluster entropy of 0.964, indicating well-balanced unit usage rather than a few dominant units. Higher K values showed progressively lower ARI and smaller minimum cluster sizes, indicating that finer inventories were not reliably recoverable from the current segment population (Figure 7, Supplementary Table S9).

The selection of K = 48 reflects a deliberate balance between vocabulary richness (enough units to capture the acoustic diversity of the corpus) and unit reliability (every unit supported by enough segment instances to be interpretable). K was not chosen to maximize downstream classification accuracy, as introducing a classification objective at the representation stage would contaminate the unsupervised acoustic discovery with label-dependent biases. The final clustering model was fit using K = 48 and fixed seed 31.

### 4.7 Token assignment and sequence construction

Each of the 4,764 segments was assigned the index of its nearest K = 48 centroid in the 128-dimensional space (units u00–u47), preserving temporal order so that each clip becomes an ordered unit sequence - e.g. a 9-segment clip might read [u27, u11, u11, u11, u11, u11, u11, u11, u27], with boundary-associated units at the ends and sustained content in between.

These sequences are the primary input to Layer 2, encoding both unit identity and the ordering and transitions between units. Mean sequence length is 8.37 (median 9, maximum 12); the 48-unit vocabulary’s 15 most frequent units account for 54% of all segment events (Figure 8).

**Figure 8.**
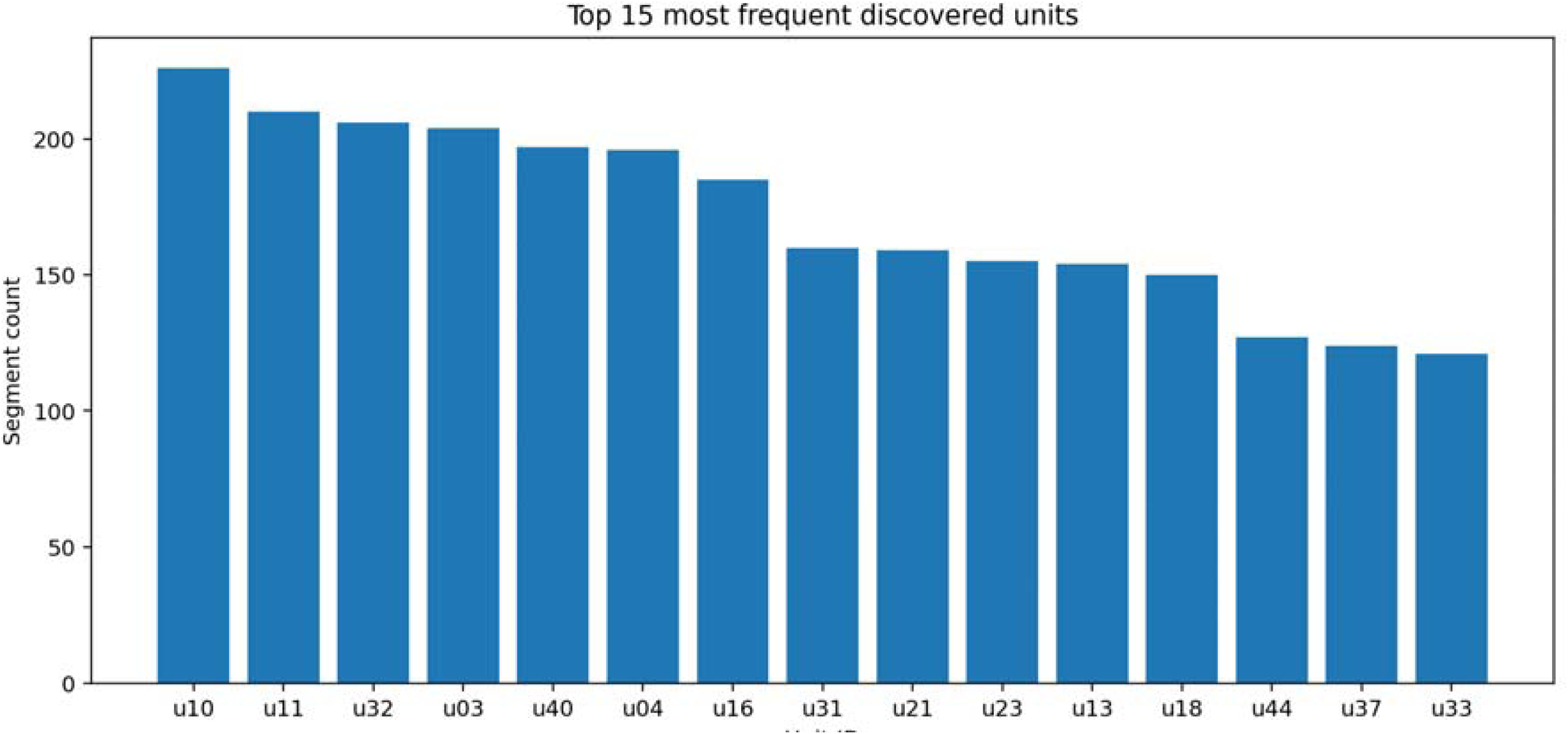
Prevalence of the 15 most frequent acoustic units in the 48-unit vocabulary, measured by segment count across all 4,764 token events. These 15 units account for approximately 54% of all segment assignments. The remaining 33 units distribute across the long tail, with no singleton clusters present in the vocabulary.

### 4.8 Dictionary scaffold construction

A dictionary scaffold linked each unit to interpretable evidence from clip metadata: three prototype segments (those nearest the centroid) as canonical instances, plus each unit’s dominant main category, subcategory, barn zone, recording context, and summary statistics (segment and clip counts, durations, global frequency). The scaffold is not a semantic dictionary - it makes no claim that a unit means a biological state - but an evidence catalog of what units co-occur with in this corpus.

### 4.9 Context bias audit

A descriptive bias audit measured whether any unit was concentrated in a single recording context, as the proportion of its segments drawn from the most frequent value of each of four variables - farm, barn zone, microphone, and microphone placement (per-unit results in Supplementary Table S6; the most context-locked units are shown in Supplementary Figure S1). A unit was flagged if its dominant proportion was ≥ 0.80 with at least 8 segments, and highly locked at ≥ 0.90 (Figure 10).

Farm identity produced the strongest locking - 9 units flagged and 6 highly locked, several with a dominant-farm proportion of 1.0 (every segment from one farm) - followed by barn zone (10 flagged, 4 highly locked); microphone and placement showed lower but meaningful concentration (Supplementary Table S10, Figure 9). The most farm-locked units (u25, u09, u27) drew all their evidence from a single farm, which precludes cross-farm inference from them and must be acknowledged in Layer 2 feature construction.

**Figure 9.**
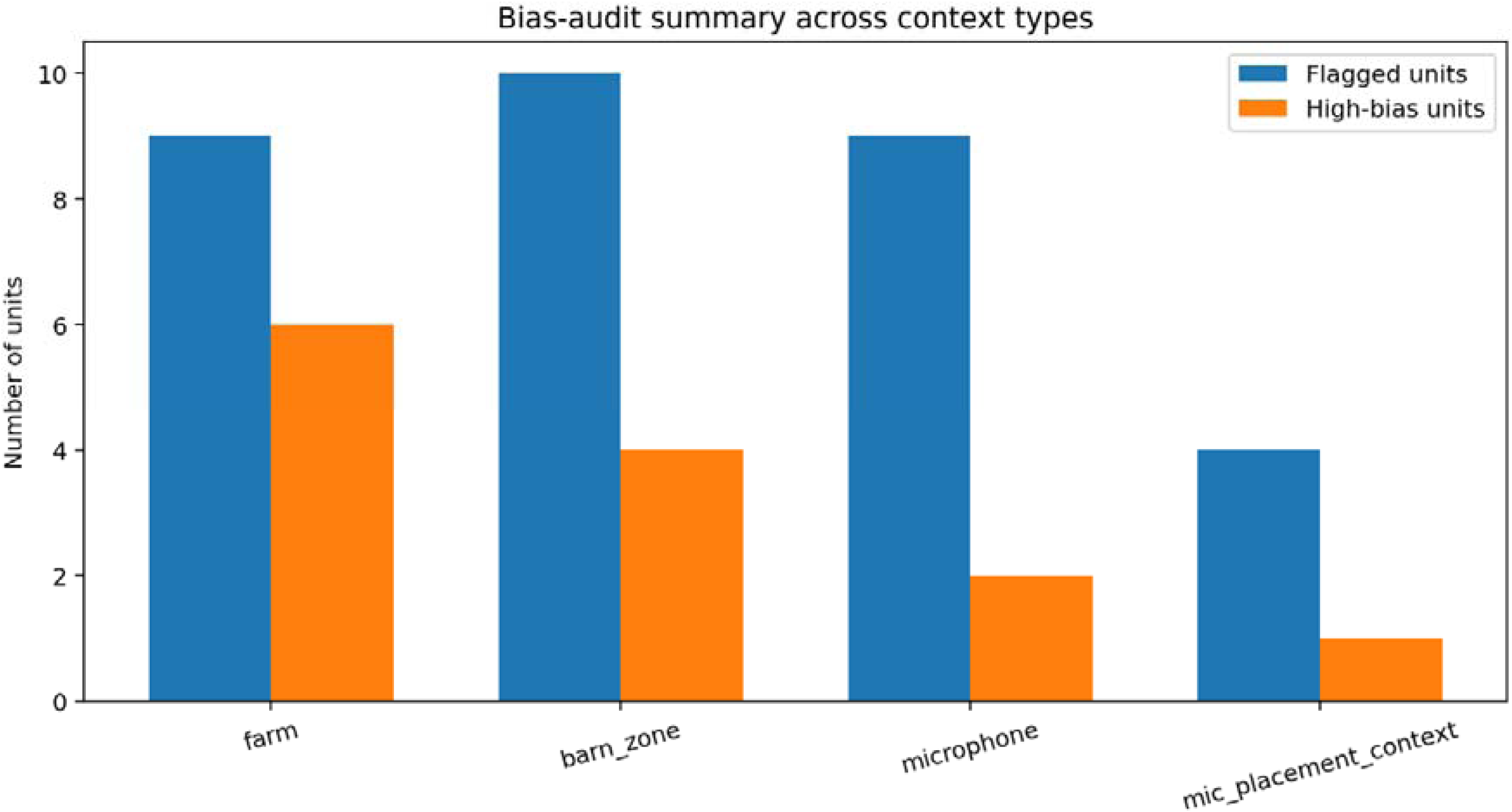
Summary of the context bias audit showing the number of flagged (dominant proportion ≥ 0.80) and highly locked (≥ 0.90) units per audited context variable. Farm identity produced the strongest locking (6 highly locked units), followed by barn zone (4 highly locked). These counts informed the construction of the high-bias-filtered vocabulary variant used in Layer 2.

The audit is descriptive, not diagnostic: a context-locked unit need not be biologically meaningless - it may reflect a phenomenon more prevalent at certain farms or zones (e.g. water-ingestion sounds at a water station). What it establishes is that such units cannot serve as farm-invariant evidence, so features built on them require either removal or transparent reporting.

The vocabulary therefore bifurcated for Layer 2 into the full 48-unit version and a high-bias-filtered version excluding the eight units highly locked across any variable (40 units), so the cost of removing the most confounded units could be measured directly.

### 4.10 Biological grounding of the unit vocabulary

The scaffold and cross-farm prototype evidence permit a preliminary biological reading of the most prevalent units, with the caveats below.

#### 4.10.1 Non-vocal physiological units (robust cross-farm group)

A small set of units dominated non-vocal physiological clips - respiration, rumination, drinking, licking - consistently across all three farms. In Farm 2 slurping and breathing prototypes a single unit occupied 9–10 of 12 positions; in a Farm 1 rumination clip a different unit filled 10 of 12 positions across 90 seconds; and a Farm 3 respiratory clip was dominated by one unit across all 12 positions. These units have longer mean durations and high global frequency, and their consistency across farms (drinking at Farm 2’s water station and Farm 1’s drinking zone, respiration at Farm 3’s resting zone) is the strongest biological grounding in the corpus, aligning with the biomechanical stereotypy of physiological sounds (Chelotti et al., 2020; Andriamandroso et al., 2016) and the cross-farm robustness of is_nonvocal (Section 7).

#### 4.10.2 Pre-milking and handling-associated units

Several units concentrated in milking-parlour recordings at Farms 1 and 3, in handling-stress, pre-milking, and novel-environment clips. Flagged as context-locked, they (e.g. u17) recur in the high-confidence failure cases of Section 7: useful as milking-context anchors but not as farm-invariant evidence.

#### 4.10.3 High-variance vocalization units

The remaining units span estrus, social, distress, maternal, and feeding calls, with higher within-clip entropy, lower global frequency, and shorter durations. The same unit appears across different behavioral categories depending on recording environment - the context-entanglement pattern of Sections 6 and 7.

#### 4.10.4 A critical caveat

These labels are inferred post-hoc from co-occurrence and prototype statistics, not validated against synchronized behavioral or physiological ground truth. The claim is not that units encode specific biological states, but that the discovery process produced a vocabulary whose patterns are consistent with the known distinction between stereotyped, farm-invariant physiological sounds and variable, context-conditioned vocalizations - qualitative support, not biological validation (Section 8).

### 4.11 Layer 1 outputs and readiness for Layer 2

At the conclusion of Layer 1, the evidence base summarized in Supplementary Table S11 was established. The token sequences, unit dictionary scaffold, and bias-audited vocabulary variants were passed directly to Layer 2 as fixed inputs. The full refined-pipeline summary metrics are reported in Supplementary Table S5. No Layer 1 parameter was re-tuned after the segmentation candidate selection was finalized, preserving the integrity of the boundary between acoustic structure discovery and proxy-state inference.

## 5. Layer 2: Context-Aware Proxy-State Inference from Discovered Token Sequences

### 5.1 Purpose and rationale

The acoustic foundation delivered a reusable symbolic representation of vocalizations as ordered token sequences, but that representation carries no biological interpretation. Layer 2 (Figure 11) tests whether these sequences carry proxy-state signal and whether contextual metadata conditions that signal in ways that survive cross-farm evaluation. Three constraints govern what can honestly be claimed: only metadata already present in the dataset were used; the targets are proxy states, not validated welfare labels, because synchronized physiological ground truth was unavailable; and evaluation spans multiple tracks so that generalization across farms, not merely predictiveness, is tested. Because dairy cow vocalizations are context-dependent signals whose meaning shifts with management, social, and physiological conditions (Briefer, 2012; Manteuffel et al., 2004), any system ignoring this dependence risks encoding farm-specific rather than general relationships. Layer 2 therefore does not decode welfare states from sound; it asks whether a label-free symbolic representation supports biologically plausible proxy inference, and how far that inference survives when an entire farm is withheld.

**Figure 10.**
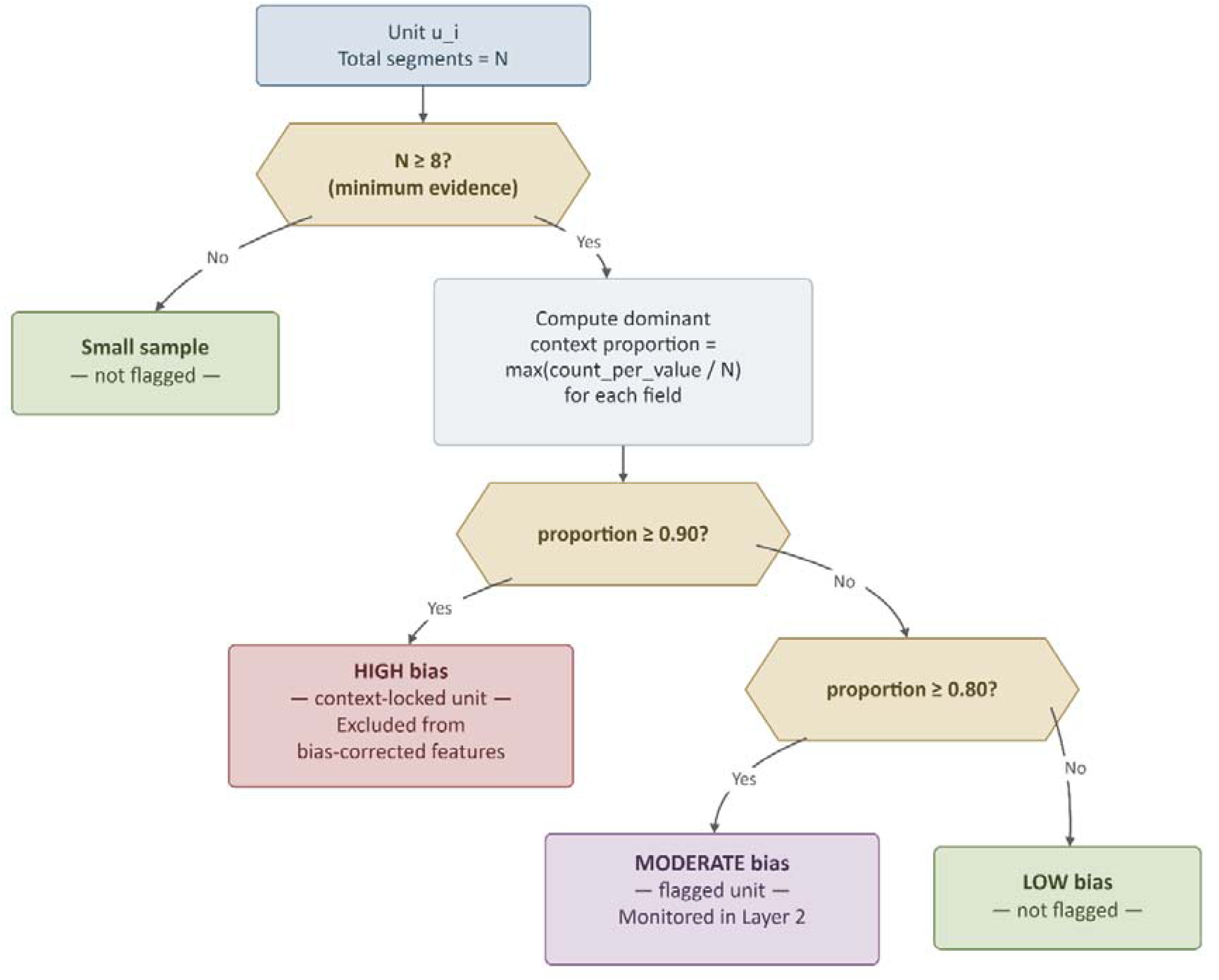
Decision logic for the context bias audit. Each of the 48 acoustic units is evaluated against four context variables (farm, barn zone, microphone, microphone placement context). A unit is flagged if its dominant context proportion reaches or exceeds 0.80 with at least 8 segment instances, and designated highly locked at 0.90. Units flagged as highly locked across any audited context variable were excluded from the bias-filtered vocabulary variant used in Layer 2.

**Figure 11.**
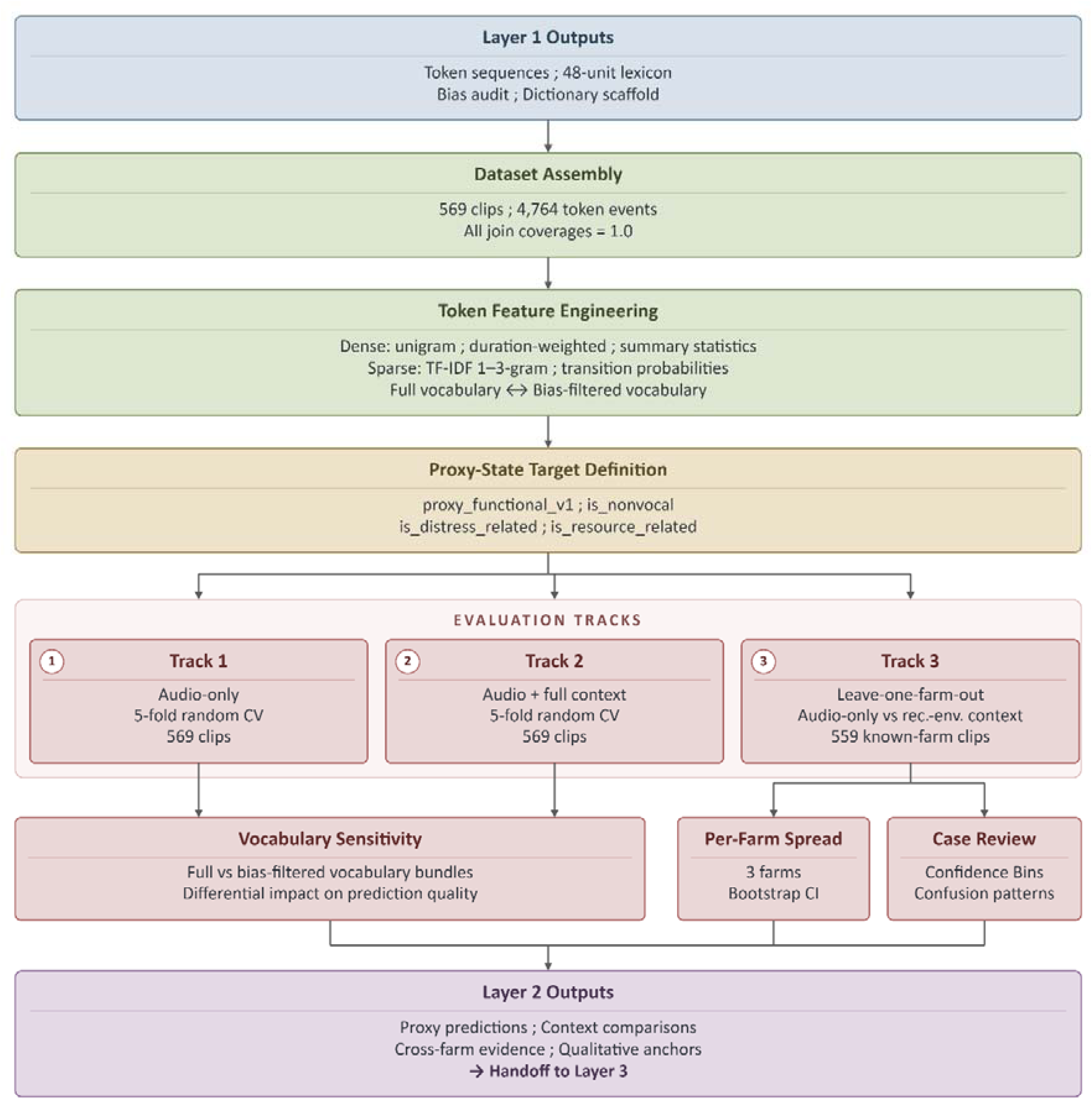
Layer 2 workflow from token-sequence feature engineering through proxy-state inference to cross-farm evaluation. Six feature families (three dense, two sparse, one summary) are computed in both full and bias-filtered vocabulary variants. Four proxy-state targets are evaluated across three matched tracks: audio-only, audio-plus-context, and leave-one-farm-out, producing per-target discriminative benchmarks and context-gain diagnostics.

### 5.2 Dataset construction and inputs from the acoustic foundation

All Layer 2 analyses were built from the refined Layer 1 branch. The clip-level master dataset was assembled by joining the manifest, the refined token sequences and token-count summaries, the unit dictionary, and the context-bias audit outputs into a single stable table. The dataset comprised 569 clips with a total of 4,764 token events and a 48-unit lexicon inherited directly from the refined branch.

### 5.3 Token-sequence feature engineering

Token sequences were transformed into six feature families. Three dense families encoded composition - unigram counts, unigram proportions, and duration-weighted proportions (scaling each token by its mean segment duration within a clip). A sequence-summary set encoded clip-level structure (sequence length, token diversity, normalized entropy, transition diversity). Two sparse families captured local order: TF-IDF-weighted 1-to-3-gram sequences, representing recurring motifs, and directed token-transition probability matrices. Each family was computed in two vocabulary variants - a full 48-unit version and a high-bias-filtered version excluding the eight units flagged as highly locked (dominant proportion ≥ 0.90) in the Layer 1 audit, leaving 40 units - yielding ten feature matrices. A 40-dimensional context block, one-hot encoding six metadata fields (farm, barn zone, microphone, microphone placement, recording day, date), was concatenated with audio features in the matched context experiments.

### 5.4 Proxy-state target definitions

Proxy-state targets mapped the original behavioral annotations into biologically plausible groupings (Figure 12). A preserved mapping (proxy_main_preserved) retained the nine original labels for reference; a compact target (proxy_functional_v1) merged them into five classes balancing biological meaning with statistical tractability; and three binary helper targets were derived for focused analysis.

**Figure 12.**
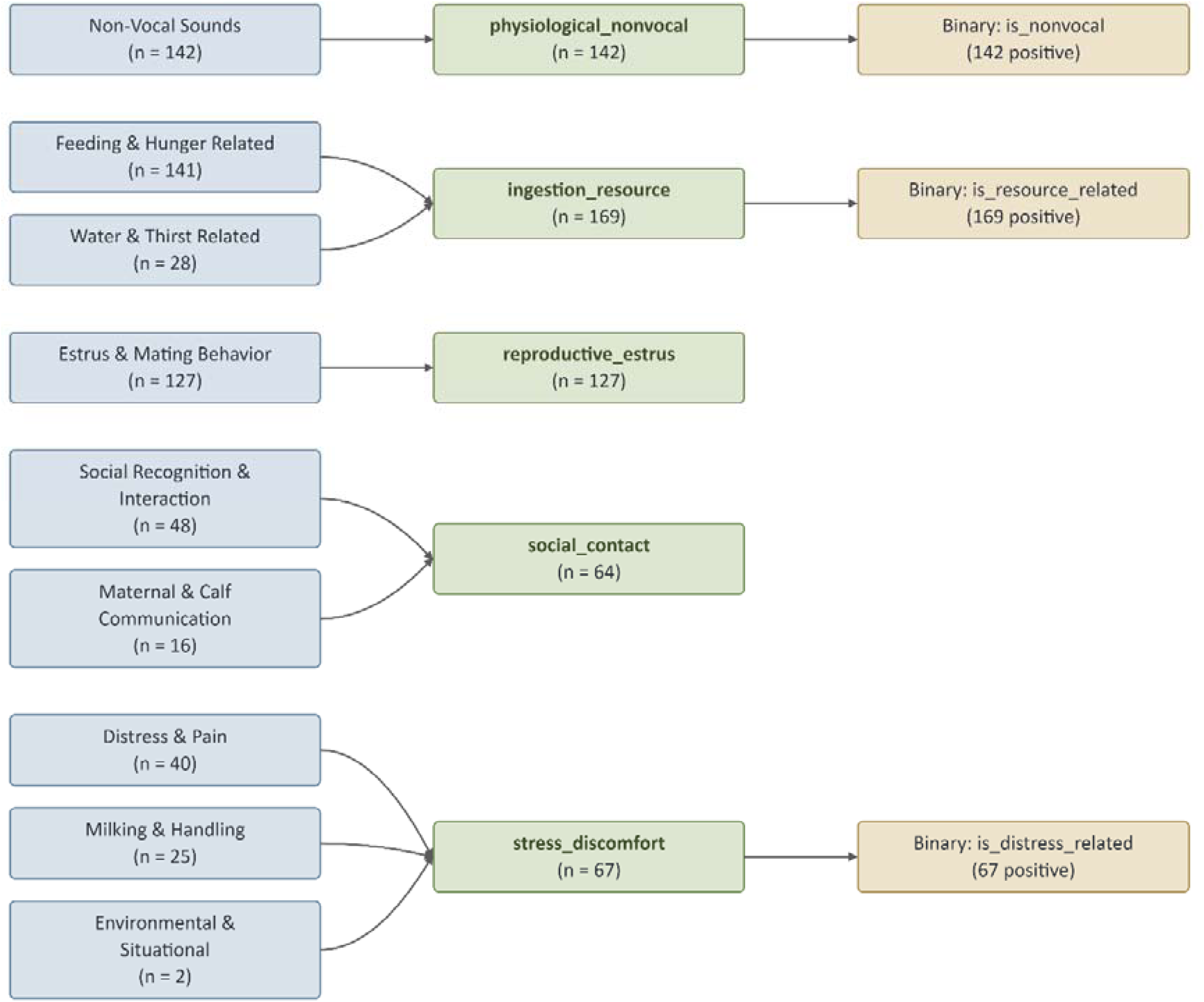
Mapping from the nine original behavioral annotation categories to the five proxy_functional_v1 classes. Merges follow shared motivational drivers: feeding and water categories form ingestion_resource; social recognition and maternal communication form social_contact; distress, handling, and environmental categories form stress_discomfort. Non-vocal sounds and estrus behavior are retained as distinct classes. Clip counts per proxy class are indicated.

The five functional classes are defined in Table 3. Physiological_nonvocal retained the non-vocal sounds intact; reproductive_estrus retained estrus and mating; ingestion_resource merged feeding and water contexts; social_contact merged social-recognition and maternal communication, whose acoustic forms overlap; and stress_discomfort merged distress and pain (n = 40), milking and handling (n = 25), and the sparse environmental and situational category (n = 2) under a shared theme of aversive or management-induced arousal. This last grouping is a conservative aversive-state proxy, not a claim of acoustic equivalence across its constituents.

These targets are proxy states rather than validated welfare labels. They rest on expert behavioral annotations made during dataset curation, not on synchronized physiological measurements or externally validated welfare outcome indicators. Their purpose is to structure biologically plausible inference questions around the token-sequence representation, not to replace welfare assessment.

### 5.5 Evaluation design

Modeling used three parallel evaluation tracks, each exposing a distinct failure mode of acoustic inference. Used together, they form a context-audited cross-farm protocol that separates acoustic generalization from farm-specific bias and makes shortcut learning empirically visible.

#### 5.5.1 Track 1: Audio-only

Five-fold stratified cross-validation on all 569 clips using token-sequence features alone, establishing the representation’s intrinsic acoustic information content.

#### 5.5.2 Track 2: Audio plus context

The same five-fold protocol with the 40-dimensional context vector concatenated to each audio matrix, testing whether context improves interpretation when farm membership is uncontrolled.

#### 5.5.3 Track 3: Leave-one-farm-out

Each of the three farms was held out in turn from the 559 known-farm clips, testing whether signal survives on an entirely unseen farm. Because farm identity as a feature would let the model absorb farm-specific shortcuts, two sub-variants were evaluated: an audio-only variant and a recording-environment variant retaining only barn zone, microphone, and placement, while excluding farm identity and temporal fields. This separates acoustic generalization from farm-identity-aided discrimination.

Within each track, the best result per target was taken from class-balanced logistic regression across all feature bundles, benchmarked against a most-frequent dummy classifier that confirmed gains reflect learned structure rather than class-frequency exploitation. For comparison, an end-to-end log-mel-spectrogram CNN was trained under the same five-fold and leave-one-farm-out protocols on the same four targets (Section 7.4, Supplementary Table S7).

### 5.6 Results

#### 5.6.1 Audio-only modeling: what token sequences carry without context

Token-sequence features produced macro-F1 values substantially above the dummy floor across all four proxy targets, establishing that the acoustic structure discovered without label supervision contains biologically plausible proxy-state signal. Supplementary Table S12 reports the best audio-only result per target.

The non-vocal physiological target produced the strongest audio-only result (macro-F1 0.808, balanced accuracy 0.846), consistent with respiratory sounds, rumination, and chewing forming a stereotyped category whose recovery needs no behavioral context. The five-class proxy_functional_v1 reached 0.654, the resource-related binary 0.704, and the distress-related binary the weakest at 0.689 - consistent with aversive-state calls being acoustically heterogeneous and context-dependent (Manteuffel et al., 2004; Briefer, 2012).

Across targets, sparse n-gram and transition features outperformed dense composition summaries, indicating that local token order adds predictive information beyond frequency - supporting treatment of the units as a structured vocabulary rather than an unordered bag.

#### 5.6.2 Audio plus available context: interpretation under matched conditions

Adding contextual metadata improved classification performance across all four targets under five-fold random cross-validation. Supplementary Table S13 reports the best audio-plus-context result per target alongside the gain over the audio-only baseline.

Gains tracked each target’s biology. The non-vocal target improved only modestly (+0.039), its signal already strong without context; the resource-related target gained most (+0.155), followed by the functional proxy (+0.112) and the distress-related target (+0.111), consistent with ambiguous states such as feeding anticipation and discomfort being more context-dependent (Supplementary Figure S2).

The best context bundles for is_distress_related and proxy_functional_v1 both used the high-bias-filtered vocabulary, suggesting that excluding the most context-locked tokens improves context-conditioning, perhaps because those tokens duplicate information already in the context block.

#### 5.6.3 Leave-one-farm-out evaluation: testing cross-farm generalization

The leave-one-farm-out analysis tested whether proxy-state signal generalized to an entirely unseen farm. Table 4 reports the best LOFO result per target together with the context variant that achieved it.

**Table 4.** Best leave-one-farm-out results by target (559 known-farm clips, 3-fold farm holdout).

| Target | Best LOFO feature family | Context variant | LOFO Macro-F1 | LOFO Bal. Acc |
| --- | --- | --- | --- | --- |
| `is_nonvocal` | TF-IDF 1–3-gram (bias-filtered) | Audio-only | 0.763 | 0.783 |
| `proxy_functional_v1` | Transition probabilities (full) | Recording-env. context | 0.407 | 0.459 |
| `is_distress_related` | Transition probabilities (bias-filtered) | Recording-env. context | 0.642 | 0.690 |
| `is_resource_related` | Unigram composition (full) | Audio-only | 0.500 | 0.500 |

The key LOFO finding is the divergence between within-cohort gains and cross-farm generalization, which was not uniform (Figure 13). The non-vocal target retained most of its within-cohort performance (LOFO macro-F1 0.763 vs 0.808), and adding recording-environment context did not help it - the acoustic signal alone is its most generalizable channel.

**Figure 13.**
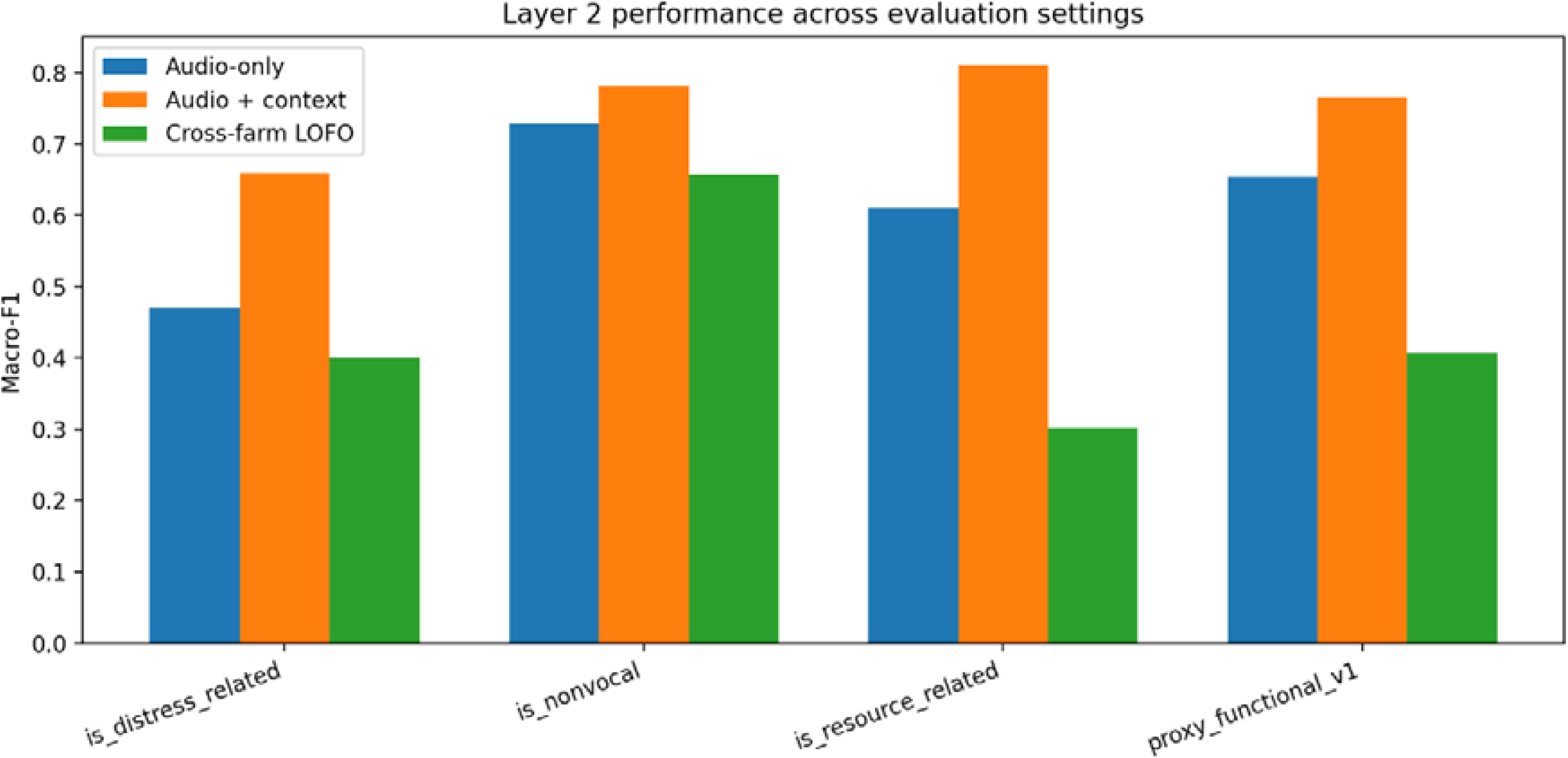
Three-track performance comparison - audio-only, audio-plus-context, and leave-one-farm-out macro-F1 - across all four proxy-state targets. The non-vocal physiological target retains the largest fraction of within-cohort performance under LOFO. The resource-related target shows the widest gap between audio-plus-context and LOFO performance, indicating strong dependence on farm-specific context shortcuts.

For the distress-related and functional targets, recording-environment context (barn zone, microphone, placement, without farm identity) gave a modest benefit, reaching 0.642 and 0.407 - well below their within-cohort audio-plus-context values of 0.800 and 0.766. Random-CV gains did not uniformly survive farm holdout.

The resource-related target had the largest within-cohort context gain (+0.155) but the weakest LOFO performance (macro-F1 0.500, indistinguishable from a majority-class baseline), and context did not help or even hurt it under holdout. The most parsimonious reading is that resource-access behavior is tightly structured by farm-specific routines, layouts, and feeding schedules, so its token patterns do not transfer between farms. This target functions as a boundary case: its collapse is not a modelling defect but a substantive finding that acoustic resource-access patterns do not generalize across commercial farms, and that performance measured without farm holdout overstates real cross-farm utility.

Across every target, adding full context including farm identity degraded LOFO performance relative to audio-only or structural context. This is direct evidence of shortcut learning: when farm identity is available, the model learns farm-specific cues that collapse on an unseen farm, so part of the random-CV context gain was farm-identifying signal rather than generalizable recording-environment information. Excluding farm identity was therefore not just a precaution but a finding - exactly the failure mode the context-audited protocol was built to surface.

#### 5.6.4 Per-farm stability and holdout spread

The farm-holdout ablation decomposed the pooled LOFO result into per-held-out-farm estimates, making the stability of cross-farm performance visible. Supplementary Table S14 reports spread statistics and bootstrap confidence intervals for the best LOFO bundle per target.

The non-vocal target was the most stable (per-farm SD 0.046; range 0.674–0.785), reinforcing its status as the most acoustically robust proxy. The functional proxy showed moderate spread (range 0.224), and the distress-related target a narrow range (0.031) driven by consistently high negative-class performance despite variable positive detection. The resource-related target’s narrow range (0.023, SD 0.010) reflects near-chance performance at all three farms (clustering around 0.459), not substantive consistency.

These statistics establish the realistic per-target performance envelope under honest farm holdout - the envelope Layer 3 must interpret (Supplementary Figure S3).

#### 5.6.5 Vocabulary sensitivity: full versus bias-filtered bundles

Comparing full and high-bias-filtered vocabularies across tracks showed only modest differences (macro-F1 gaps generally below 0.015 under random CV); the filtered vocabulary was slightly preferred for the non-vocal and distress targets under LOFO, the full vocabulary for the resource and multiclass targets (Supplementary Figure S5). Predictive signal is thus not concentrated in the most context-locked tokens: excluding them neither systematically helps nor harms. Context-locking is a second-order source of variation relative to farm-level differences, and both variants were carried into Layer 3.

#### 5.6.6 Qualitative case review and confusion structure

A qualitative review was conducted over the best LOFO bundle per target across 559 known-farm clips. Supplementary Table S15 summarizes descriptive accuracy and confidence statistics from this review.

Two confidence patterns matter for Layer 3. For the non-vocal and distress-related targets, the model was more confident when wrong (0.566, 0.542) than when right (0.352, 0.262) - errors are high-confidence, not cautious - so calibration and abstention are especially relevant. For proxy_functional_v1 the relationship was healthy: confidence was higher for correct (0.654) than incorrect (0.442) cases, so confident multiclass predictions are a relatively reliable guide to correctness.

In the functional proxy, ingestion_resource was the most error-prone class, most often confused with social_contact (51), stress_discomfort (43), and physiological_nonvocal (40), with Farm 2 contributing most of these errors when held out - indicating that resource-related token patterns at Farm 2 overlap with other categories’ signatures (Supplementary Figure S4). Reproductive_estrus and social_contact were better separated. The problem is therefore not uniformly hard: the most confused boundaries are those with the greatest acoustic overlap under farm-specific context.

### 5.7 Synthesis

Layer 2 established three findings. Discovered token sequences carry proxy-state signal without any context; available context improves interpretation under within-cohort evaluation, especially for ambiguous targets; and cross-farm robustness is selective - the non-vocal target held up under farm holdout (LOFO macro-F1 0.763) while the resource-related target fell to chance (0.500), with the distress-related and multiclass targets intermediate. This selective robustness is itself informative: it identifies which proxy categories are acoustically recoverable across farms and which need farm-specific context to be useful. It also frames the next layer’s question - not whether these predictions contain signal (they do) but whether the probabilities attached to them are trustworthy enough for confidence-aware interpretation, and when the system should abstain rather than predict.

## 6. Layer 3: Reliability, Calibration, Selective Prediction, and Model-Behaviour Dictionary

### 6.1 Purpose and role

Layer 3 does not alter the upstream acoustic representation, does not re-open feature engineering, and does not re-label any data. Its purpose is narrower but scientifically essential: to determine how trustworthy the proxy-state probabilities produced in Layer 2 are, under what conditions the system should abstain rather than predict, and how the discovered acoustic units can be summarized into a structured evidence lexicon without overclaiming semantic certainty.

The distinction between classification accuracy and probability quality is central to this layer. A model can produce correct hard-label predictions while assigning miscalibrated confidence values, and a model with well-calibrated probabilities may not always maximize macro-F1. Layer 3 therefore evaluates these two properties separately. It treats calibration as a post-processing selection problem solved independently for each target, asks whether selective prediction on uncertain clips increases the reliability of retained outputs, and constructs a model behaviour token-state dictionary whose entries represent evidence tendencies rather than deterministic mappings. Together, these steps convert the Layer 2 output from a set of predictions into a calibrated, abstention-aware evidence layer. Figure 14 presents the overall Layer 3 architecture.

**Figure 14.**
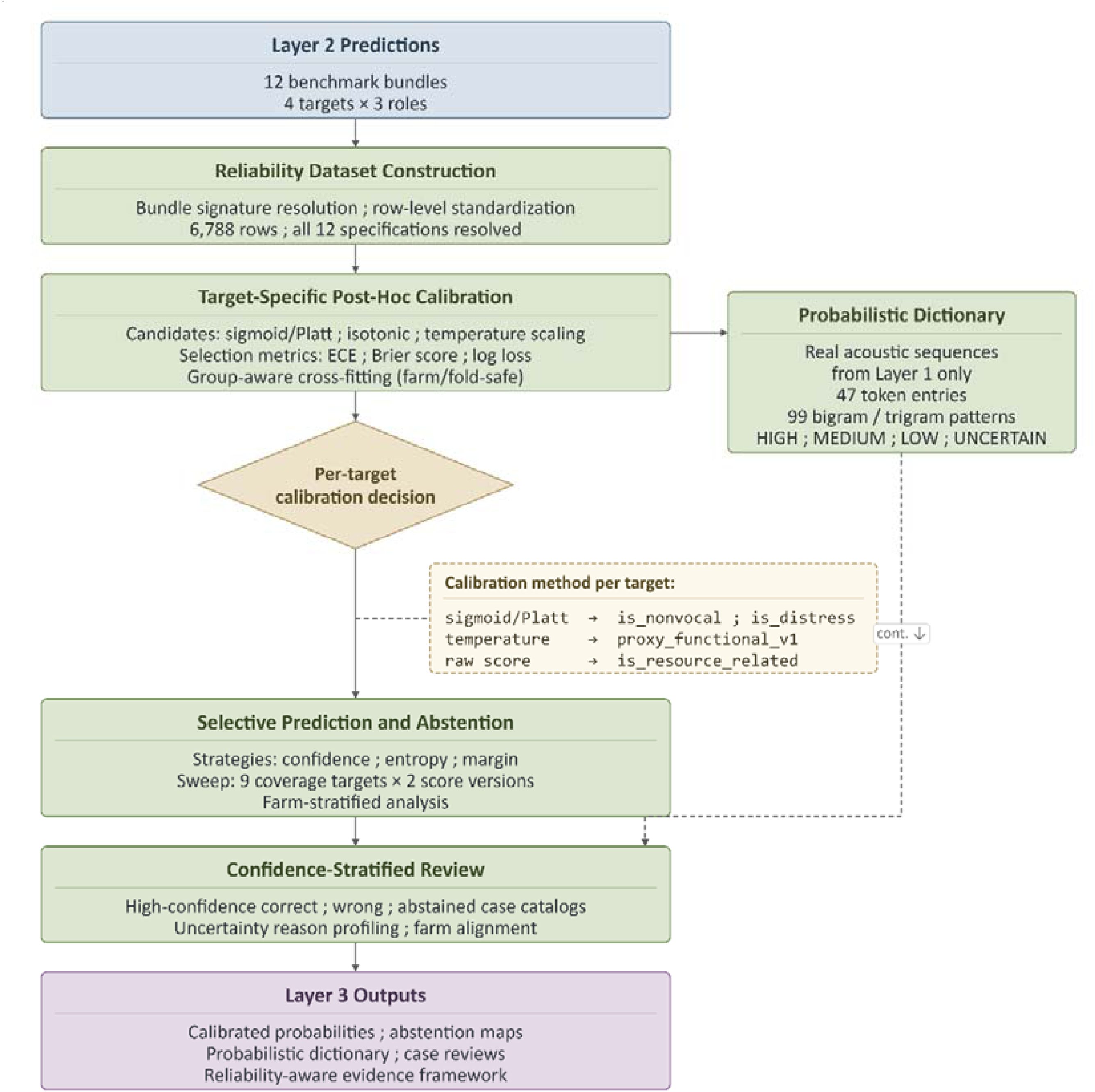
Layer 3 architecture overview. The reliability assessment pipeline receives fixed benchmark bundles from Layer 2 and proceeds through four stages: target-specific post-hoc calibration selection, selective prediction under confidence-based abstention, model-behaviour dictionary construction with reliability-tier assignment, and confidence-stratified clip-level review. Each stage produces auditable outputs that feed into the final reliability summary.

### 6.2 Reliability dataset construction

Layer 3 first assembled a unified reliability dataset from the Layer 2 outputs. Each benchmark bundle was resolved by feature signature - feature family, vocabulary variant, context variant, model, and evaluation track - rather than by ad hoc row selection, so that calibration never operated on misidentified outputs. The dataset spanned four targets × three benchmark roles per target: the audio-only reference, the best random-CV context reference, and the leave-one-farm-out benchmark (the primary track for Layer 3). It contained 6,788 rows - 559 per LOFO bundle and 569 per random-CV bundle - covering all 569 clips with valid probability dictionaries, with class distributions matching those reported in Layer 2. The three-role structure was preserved throughout so that every calibration and reliability result remained tied to its correct evaluation context.

### 6.3 Target-specific post-hoc calibration

#### 6.3.1 Calibration design and rationale

The calibration analysis in Layer 3 addressed a known limitation of applying a single post-hoc calibration strategy uniformly across all targets. Different proxy-state targets differ in class structure, class imbalance, farm sensitivity, and the geometry of their raw probability distributions. A single calibration recipe that improves one target may degrade another, particularly when the binary and multiclass settings require different calibration machinery.

A target-by-target selection approach was therefore adopted. For each target, calibration candidates appropriate to its class structure - raw probabilities, Platt/sigmoid scaling, isotonic regression, and temperature scaling for the binary targets; raw, multiclass temperature scaling, and one-vs-rest Platt and isotonic for the multiclass target - were evaluated on the primary LOFO bundle and ranked on probability quality alone (expected calibration error, Brier score, and log loss), never on accuracy or macro-F1. Calibrators were fit in a group-aware manner, using only the remaining farms when recalibrating a held-out farm, so that the held-out farm never appeared in the calibrator’s training set. The full candidate sets and selection safeguards are detailed in Supplementary Methods S3; the per-target selection logic is shown in Figure 15.

**Figure 15.**
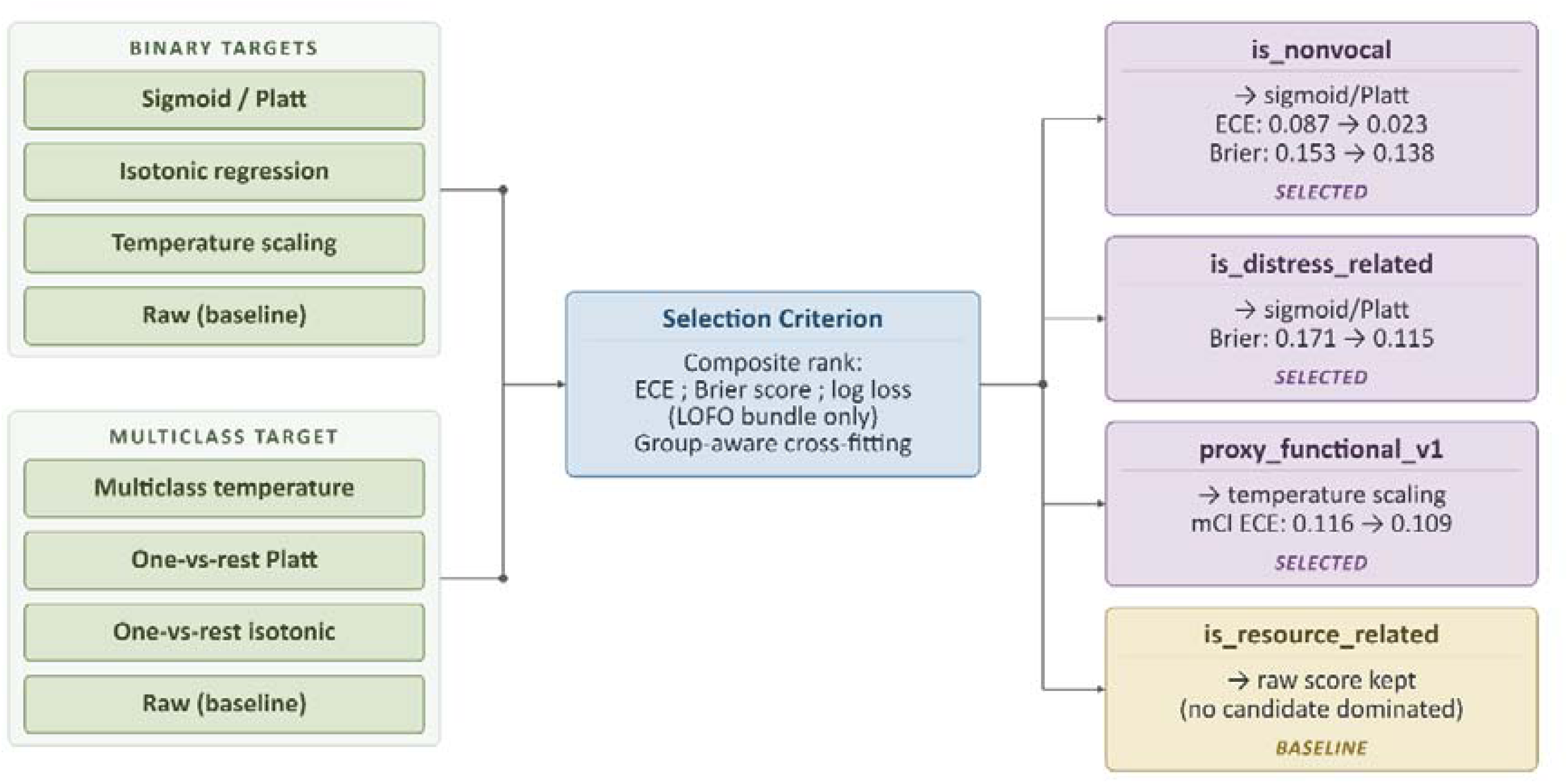
Calibration selection logic applied independently to each proxy-state target. Binary targets are evaluated against raw, sigmoid/Platt, isotonic, and temperature scaling candidates; multiclass targets against raw, temperature scaling, one-vs-rest Platt, and one-vs-rest isotonic. Selection is made on the primary LOFO bundle using composite probability-quality ranking (ECE, Brier score, log loss). The selected method per target is indicated at right.

#### 6.3.2 Calibration outcomes

For the non-vocal physiological target, calibration provided the clearest and most consistent improvement (Table 5; Supplementary Figure S7). Expected calibration error fell from 0.087 to 0.023, a reduction that substantially narrowed the gap between predicted confidence and empirical accuracy. This improvement was achieved without meaningful sacrifice in Brier score, and the sigmoid/Platt method was ranked first across all composite metrics. The non-vocal target was also the most interpretable calibration benchmark in this study, because its acoustic structure is relatively stereotyped and its raw predictions were already among the most confident.

**Table 5.**
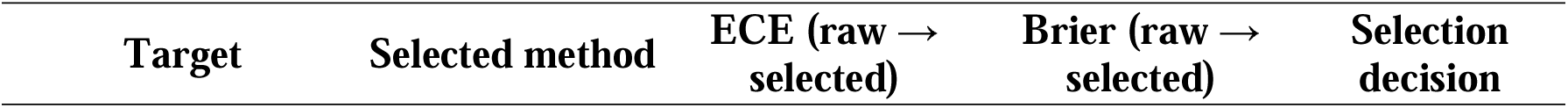
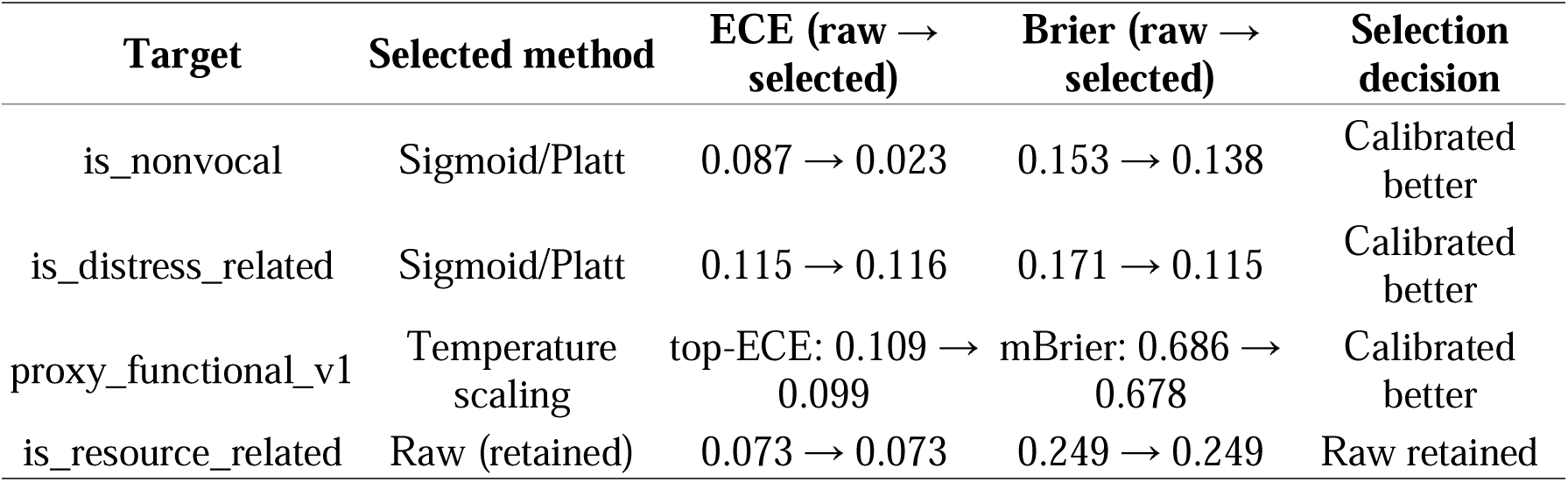
Post-hoc calibration selection and probability-quality outcomes by target (primary LOFO bundle).

For the distress-related target the result was more nuanced. Sigmoid/Platt scaling improved the Brier score (0.171 to 0.115) while top-label ECE was essentially unchanged (0.115 to 0.116) - a known behaviour when calibrating an imbalanced binary target toward its base rate, since ECE is measured on the predicted argmax class. The composite ranking still selected sigmoid/Platt. Because this recalibration also shifted the decision boundary toward the majority class, calibrated macro-F1 (0.483) fell below the raw value (0.642); this is the expected cost of honest probabilities, not a failure of calibration. Discriminative comparisons across the pipeline should therefore use the raw LOFO macro-F1 from Layer 2, and probability-quality metrics the calibrated values.

For proxy_functional_v1, temperature scaling provided modest but consistent improvements across both multiclass Brier and top-label ECE. The temperature values fitted per fold ranged from 0.65 to 1.90, reflecting heterogeneity across the three farm holdout splits. This heterogeneity itself is informative: a well-calibrated multiclass model should require similar temperature values across all folds, and the variability here signals that the proxy_functional_v1 probability geometry differs substantially from one farm to another.

For the resource-related target, no calibration candidate improved upon raw probabilities on the primary LOFO bundle under the composite ranking. Raw probabilities were therefore retained. Preserving raw probabilities for is_resource_related means the Layer 3 analysis correctly identifies that this target’s probability structure is already at or near the plateau of what simple post-hoc calibration can achieve for this dataset, not that calibration was not attempted.

### 6.4 Selective prediction and abstention

#### 6.4.1 Design and motivation

Even well-calibrated probabilities do not guarantee that every prediction is equally trustworthy. For clips where the model is uncertain - where predicted probabilities are spread across classes, where confidence margins are narrow, or where the token sequence contains many context-locked units - producing a hard prediction may be less informative than acknowledging uncertainty. Selective prediction evaluates whether withholding low-confidence predictions improves the quality of the retained subset and at what coverage cost.

Three abstention strategies were evaluated: confidence-based abstention (withholding clips whose maximum predicted probability falls below a threshold τ), entropy-based abstention (withholding clips with high predictive entropy), and margin-based abstention (withholding clips with small top-2 probability gaps). Each strategy was evaluated on both raw and calibrated probability scores. Coverage targets of 100%, 95%, 90%, 85%, 80%, 75%, 70%, 60%, and 50% were tested systematically. Farm-stratified abstention summaries were also computed to expose whether uncertainty clustered within specific farms.

A practical operating point at 80% coverage was used as the primary summary point, supplementing a fixed-threshold analysis at τ = 0.70. The 80% coverage operating point was chosen to represent a deployment scenario where approximately one in five predictions is deferred to a human observer, which is a realistic operating regime for advisory monitoring tools in precision livestock farming.

#### 6.4.2 Selective prediction outcomes

The abstention results (shown in Table 6) differed substantially across targets, and the pattern of results was scientifically informative rather than uniformly positive (Supplementary Figure S8).

**Table 6.** Selective prediction at the 80% coverage operating point by proxy-state target (primary LOFO bundle, selected score version).

| Target | Score version | Baseline macro-F1 | Retained macro-F1 (80% cov.) | Abstention rate | Interpretation |
| --- | --- | --- | --- | --- | --- |
| is_nonvocal | Raw | 0.763 | 0.793 | 10.2% | Modest selective gain |
| proxy_functional_v1 | Calibrated | 0.407 | 0.430 | 14.0% | Meaningful selective gain |
| is_distress_related | Calibrated | 0.483 | 0.483 | 0.0% | No selective benefit |
| is_resource_related | Raw | 0.500 | 0.503 | 7.9% | Negligible selective benefit |

For the non-vocal target, raw confidence was a useful abstention signal: retained macro-F1 rose from 0.763 to 0.793 at 80% coverage, and to 0.807 at τ = 0.70 (withholding 43.5% of clips). Calibrated confidence was not useful here - sigmoid/Platt scaling made it a poorer rank-ordering of correctness than raw confidence - illustrating that calibration (aggregate probability quality) and abstention (confidence rank-ordering by correctness) are different objectives that can diverge.

For proxy_functional_v1, abstention was beneficial under calibrated scores: retained macro-F1 rose from 0.407 to 0.430 at 80% coverage, the temperature-calibrated confidence reliably identifying a subset of trustworthy predictions.

For the distress-related target, abstention gave no retained-set benefit under calibrated scores (only 3.4% withheld at τ = 0.70, macro-F1 unchanged at 0.483); a modest raw-score gain of 0.026 was possible at 68.7% coverage. Calibration toward the distress base rate left little confidence variation to exploit - better-calibrated probabilities but a weaker abstention signal.

For the resource-related target, abstention provided negligible benefit at any threshold, consistent with its status as a boundary case: raw probabilities were retained and selective prediction could not rescue cross-farm performance.

### 6.5 Model-behaviour dictionary construction

#### 6.5.1 Design and framing

The model-behaviour dictionary extends the Layer 2 acoustic-unit framework into an interpretable evidence lexicon: for each discovered acoustic unit, it records how that unit’s presence in a clip’s token sequence tends to associate with proxy-state probability profiles under the Layer 3 reliability framework. The intent is an interpretable intermediate representation between the discovered acoustic units and downstream decision support, reflecting within-corpus model behaviour under calibrated conditions rather than validated biological meaning. A token’s entry therefore carries a bounded claim: within the current corpus and under the current proxy-state targets, clips containing that token tended to receive certain probability profiles and certain reliability characterizations.

The dictionary was built only from the real Layer 1 token sequences and resolves 47 of the 48 discovered units; one unit was absent from all LOFO evaluation sequences and so received no profile. Each token was assigned a reliability tier per target - HIGH, MEDIUM, LOW, or UNCERTAIN - from three jointly applied criteria: its mean confidence across clips containing it, the fraction of those clips flagged uncertain, and its evidence count. The tier thresholds were target-specific, reflecting the differing probability scales of the four targets (Supplementary Methods S4). These tiers describe the within-corpus reliability of a token’s evidence profile; they do not describe whether the token generalizes across farms.

#### 6.5.2 Tier distribution and interpretation

The tier distribution for is_nonvocal is the most differentiated (Figure 16), with tokens spread across all four tiers and the largest proportion in the UNCERTAIN category. This reflects a spread in how consistently different acoustic units associate with the non-vocal proxy state: some units are strongly and consistently associated (HIGH), while others show inconsistent or low-confidence associations (UNCERTAIN or LOW). The five polysemous candidates for this target - tokens for which predictive entropy was high and the dominant predicted class was weak - indicate acoustic units whose token-sequence context changes the implied state.

**Figure 16.**
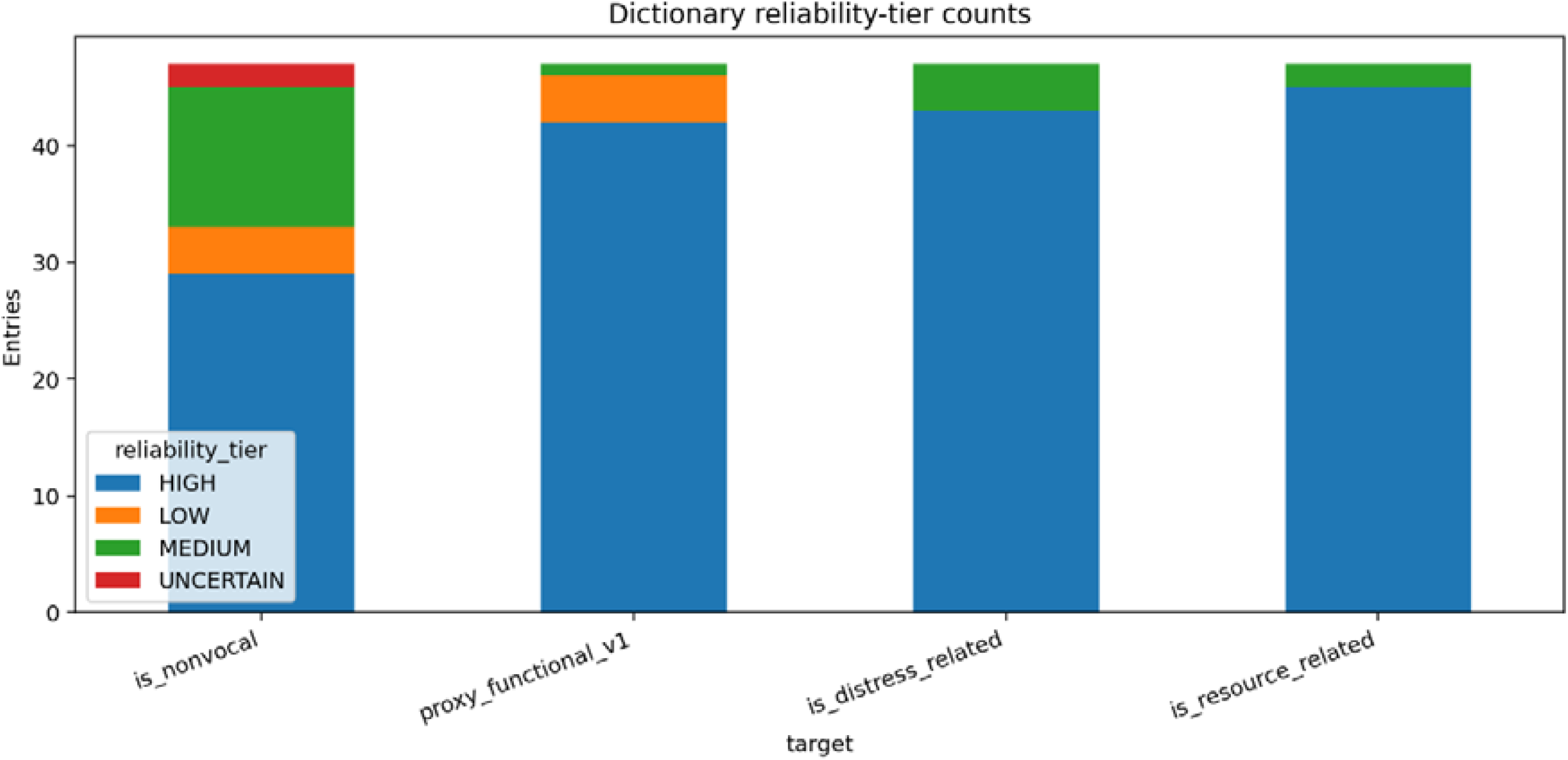
Reliability tier distribution across all four proxy-state targets in the model-behaviour dictionary (47 profiled tokens per target). The is_nonvocal target shows the most differentiated spread across HIGH, MEDIUM, LOW, and UNCERTAIN tiers. The distress-related and resource-related targets concentrate the majority of tokens in the HIGH tier, reflecting within-corpus confidence patterns that do not correspond to cross-farm discriminative strength.

The tier distributions for is_distress_related and is_resource_related place most tokens in the HIGH tier, at 91.5% and 95.7% respectively (Supplementary Table S16). These numbers require explicit qualification. The HIGH tier reflects within-corpus confidence distributions calibrated to the target’s raw threshold, not cross-farm generalization. For is_distress_related, the raw LOFO macro-F1 was 0.642, and for is_resource_related it was 0.500 at balanced accuracy 0.500. These cross-farm performance figures are entirely consistent with high within-corpus confidence: a model can consistently assign high confidence to the majority class (non-distress, non-resource) and thereby exhibit low ECE and high mean confidence while failing to generalize its minority-class distinctions across farms. The dictionary entries for these two targets should therefore be interpreted as evidence profiles reflecting within-corpus model behavior, not as cross-farm validated evidence of the acoustic units’ biological relevance to those proxy states. The stress-test designation of is_resource_related applies to the dictionary as well as to the modeling results.

Polysemous candidates were flagged across all four targets, totalling 7 for proxy_functional_v1 alone. These are acoustic units for which predictive entropy exceeded the 85th percentile and the dominant predicted class accounted for fewer than 60% of predictions. They represent tokens whose association with proxy states appears context-dependent - the same token, in different clip contexts, aligns with different proxy tendencies - and they support the Layer 2 finding that token meaning is partly context-conditioned.

### 6.6 Confidence-stratified review

The Layer 3 reliability analysis operates on the same primary LOFO benchmark bundles identified as best-performing in Section 5.6.3, resolved by feature-bundle signature in the dataset construction step (Section 6.2). The confidence statistics reported below were computed through the Layer 3 probability analysis pipeline and differ from the qualitative confidence values in Section 5.6.6, which were generated through a separate review pipeline with a different confidence definition. The confidence-stratified review examined the relationship between prediction confidence and correctness at the clip level, drawing on the 559 known-farm LOFO clips across all four targets. The review was constructed from the clip-evidence table and all fragile derived fields were recomputed from scratch rather than inherited from upstream flags.

Two confidence patterns are notable (Figure 17). For proxy_functional_v1, the mean calibrated confidence (0.414) is lower than the mean raw confidence (0.535) (Supplementary Table S17), because temperature scaling with values above 1.0 softens probability distributions and reduces their peak. This is the intended behavior of temperature scaling under over-confident raw predictions, and it is consistent with the calibration improvement in top-label ECE from 0.109 to 0.099. For is_resource_related, raw and calibrated confidence are identical at 0.633, confirming that raw probabilities were retained without modification.

**Figure 17.**
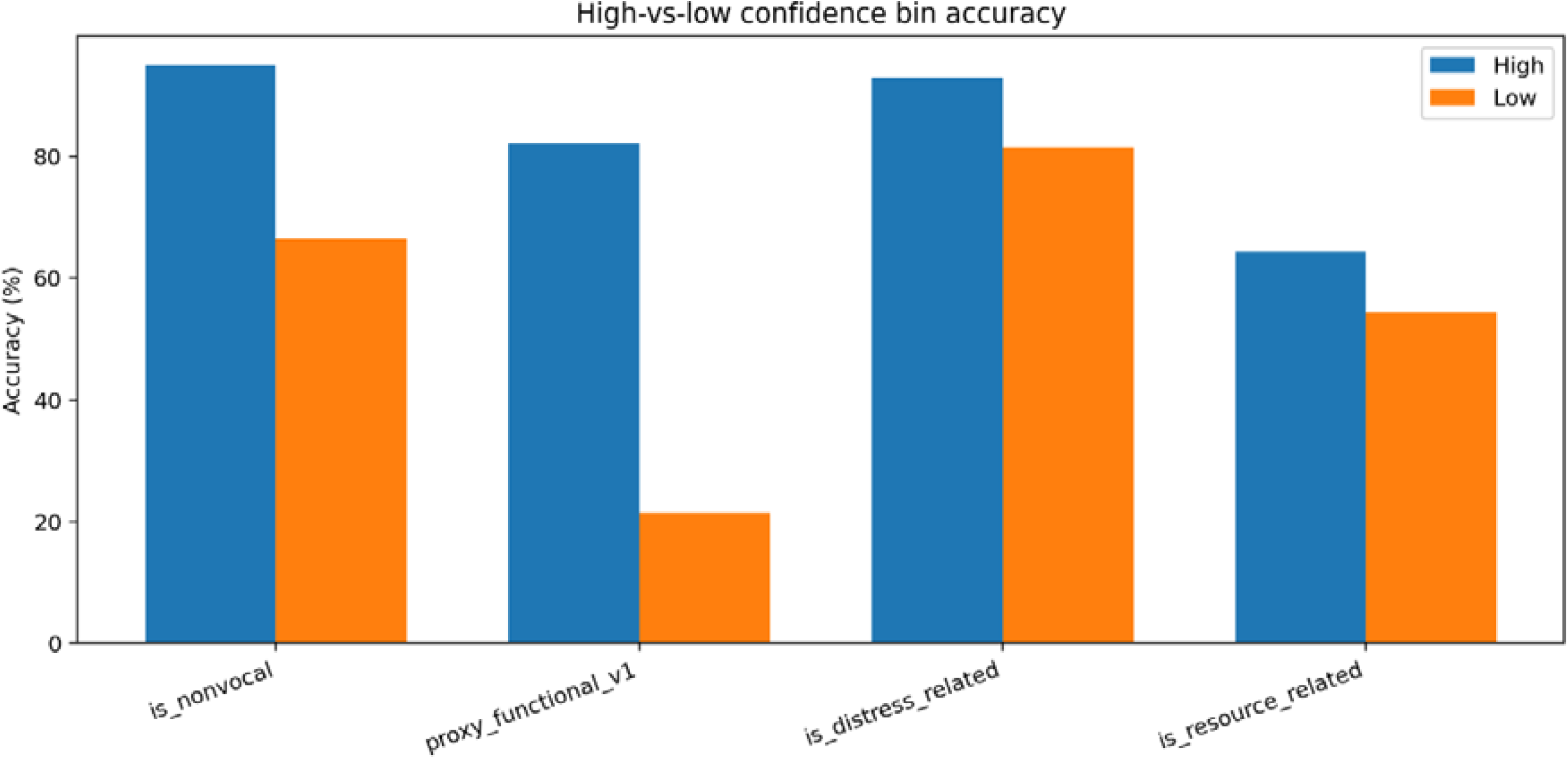
Relationship between model confidence and empirical accuracy across the four proxy-state targets. Each target is shown with high-confidence and low-confidence strata, revealing the calibration structure of the primary LOFO benchmark.

The uncertainty reason analysis (shown in Figure 18) identified the most frequent structural sources of prediction uncertainty across all four targets. Token-sequence factors were dominant: repeated-token dominance within a clip (166 clips per target), low within-clip token diversity (150 clips), high predictive entropy (140 clips), and low raw confidence (140 clips) each appeared consistently across all targets and all three farms. These reasons are not mutually exclusive - a single clip may trigger several simultaneously - but their consistent appearance across farms and targets suggests that within-clip sequence structure, not farm-specific recording conditions alone, is a primary driver of model uncertainty.

**Figure 18.**
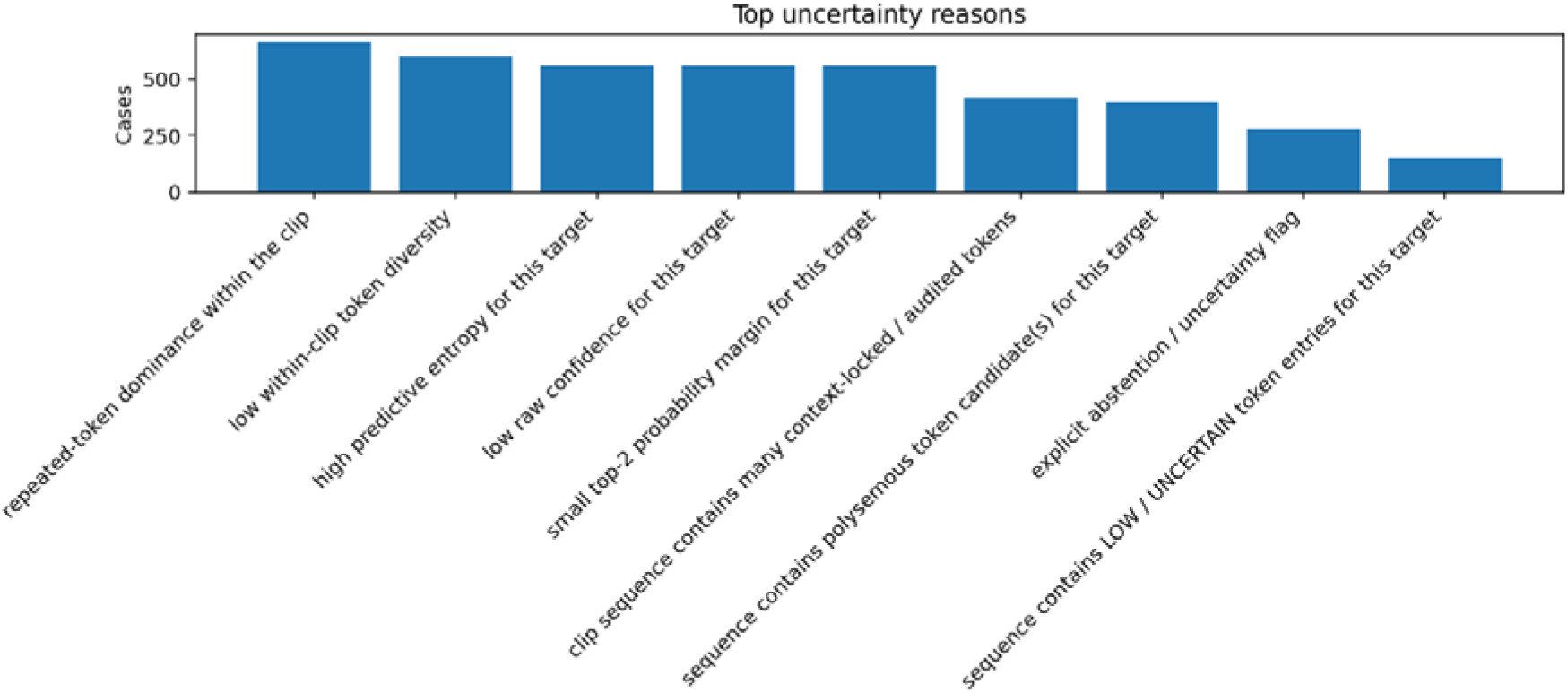
Most frequent structural uncertainty reasons identified across all four proxy-state targets in the confidence-stratified review. Token-sequence factors - repeated-token dominance, low within-clip diversity, high predictive entropy, and low raw confidence - appear consistently across targets and farms, indicating that within-clip sequence structure is a primary driver of model uncertainty independent of recording site.

The farm-level accuracy patterns reinforced the Layer 2 finding that performance varies across farms, particularly for the weaker targets. For is_resource_related, per-farm accuracy ranged from 0.453 to 0.758, with the worst farm (Farm 2) falling well below the aggregate. For is_nonvocal, per-farm accuracy ranged from 0.767 to 0.851, a substantially narrower spread (Supplementary Figure S6). The confidence-stratified review thus translates the quantitative cross-farm spread statistics from Layer 2 into a clip-level inspection framework, making the sources of uncertainty visible at the instance level rather than only at the aggregate level.

### 6.7 Layer 3 synthesis

Layer 3 achieved what it was designed to achieve: it distinguished between targets where reliability-aware analysis adds value and targets where it reveals honest limits. The non-vocal physiological target benefits from post-hoc calibration, improving its ECE by 0.064 (Supplementary Table S18), and its raw confidence scores support meaningful selective prediction. The multiclass proxy target benefits modestly from both temperature calibration and abstention at practical coverage levels. The distress-related target shows improved Brier score after calibration, but the calibrated model’s decision boundary collapses toward majority-class prediction, and abstention provides no retained-set benefit under calibrated scores. The resource-related target receives no benefit from any Layer 3 procedure, correctly marking it as the boundary case of the current framework.

The three layers form a coherent evidence-building chain. The resulting framework does not produce a single scalar welfare score. It produces a differentiated reliability map - some targets more trustworthy, some less, some benefiting from abstention, some not - which is the appropriate output of a system that takes scientific honesty about its own limitations seriously.

## 7. Discussion

### 7.1 What the pipeline achieved

The central finding of this study is that unsupervised acoustic unit discovery, combined with context-aware proxy-state inference and reliability-aware interpretation, can produce a differentiated and defensible evidence framework without synchronized welfare ground truth. The contribution is not a new learning algorithm but the integration and evaluation logic: a representation discovered without a fixed label space, a farm-held-out protocol that exposes shortcut learning, and probability estimates made more honest through calibration. This departs from the supervised, single-site classification designs that dominate cattle vocalization analysis (Gavojdian et al., 2024; Meen et al., 2015), which treat classification accuracy as the primary evidence of meaning. Table 7 summarises each target’s trajectory from audio-only signal through cross-farm evaluation to Layer 3 reliability, and the interpretive label it earns.

**Table 7.**
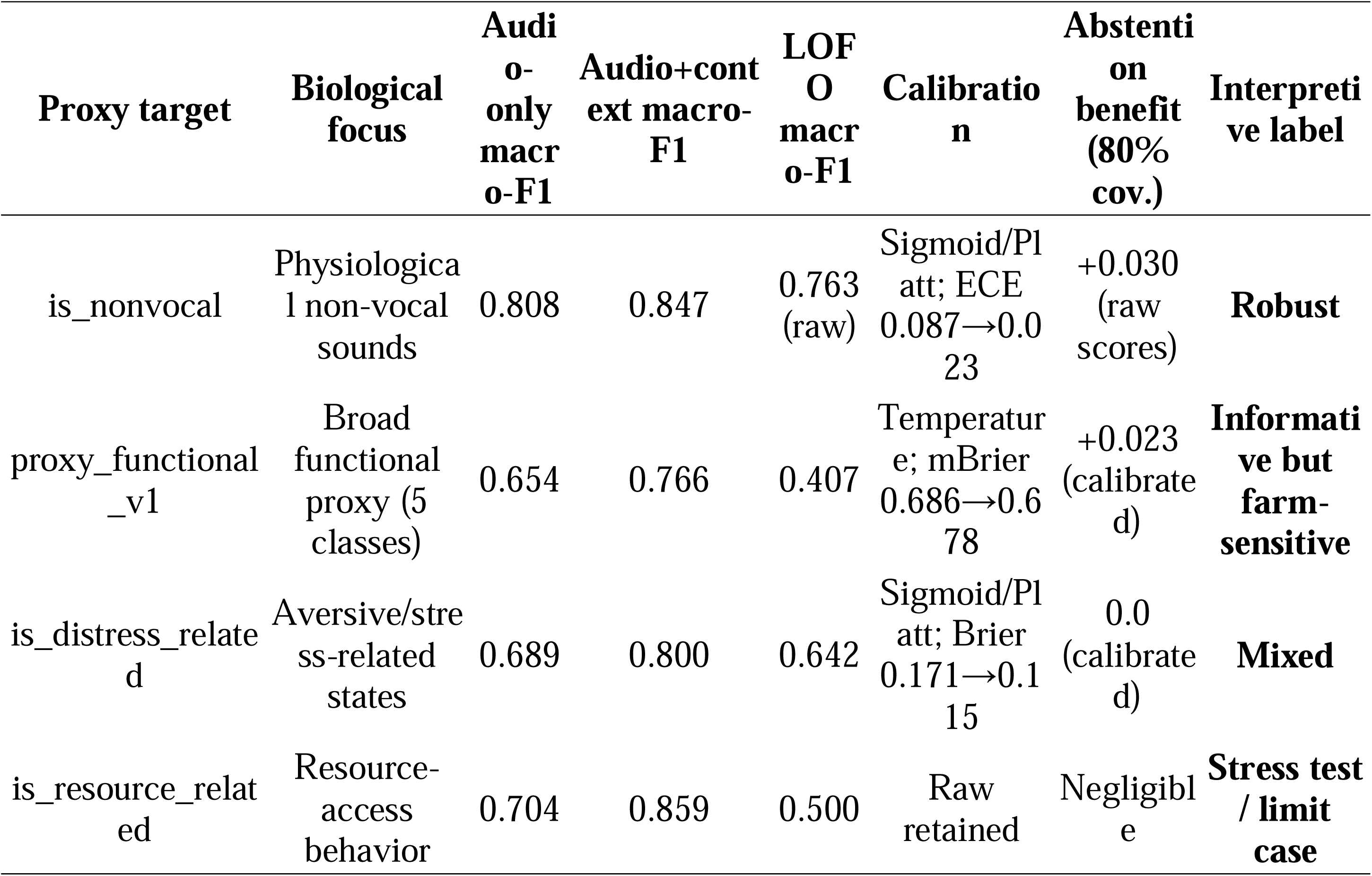
Master evidence summary across all three analytical layers.

The span of that summary - from a robust cross-farm result with calibrated ECE of 0.023 to a stress-test target that gained nothing from any reliability procedure - is itself a result: vocalization-based proxy inference is not uniformly tractable, and its tractability is biologically structured rather than random.

### 7.2 Why physiological non-vocal structure is robust across farms

The cross-farm generalizability of the non-vocal physiological proxy is the study’s most important single finding, and it is not obvious why an unsupervised representation should transfer across farms with different housing, microphones, and layouts. The explanation is biomechanical. Respiratory events, rumination, chewing, drinking, and licking are produced by stereotyped motor programs whose acoustic signatures are set by the animal’s anatomy rather than its motivational state (Chelotti et al., 2020; Andriamandroso et al., 2016). Unlike communicative vocalizations, which encode internal state and are sensitive to social and management context (Briefer, 2012), these sounds are by-products of physiology; their aperiodic, broadband, turbulence-dominated structure is largely farm-invariant. The token evidence supports this: units associated with drinking and slurping dominated clips from Farms 1 and 2, a short respiratory unit dominated a sneezing clip from Farm 3, and a single repeated unit dominated rumination recordings - whereas the same vocabulary applied to estrus, social, and hunger calls produced diverse, variable sequences. This contrast between biomechanically determined sounds and communicatively flexible calls explains why one proxy generalizes and others do not, and suggests that systems separating physiological monitoring from behavioral vocalization analysis may generalize better than those treating all cow sounds in one classifier (Jobarteh et al., 2025; Kate and Neethirajan, 2026).

### 7.3 Why context helps some targets but limits cross-farm generalization

Contextual metadata produced a consistent but counterintuitive pattern. Under random cross-validation, adding context improved all four targets, with the resource-related proxy gaining most (+0.155 macro-F1); under leave-one-farm-out, those gains did not survive for the weaker targets, and recording-environment context sometimes degraded performance relative to audio alone. The explanation is context entanglement. Resource- and distress-related states have acoustic expressions shaped by feeding routine, pen layout, and spatial setting: a cow calling near an empty feeder at one farm does not sound like a cow in the same state at another, where feeding systems, barn geometry, and microphone placement differ. When farm identity is available - directly or through correlated variables such as barn zone and microphone placement - the model learns these farm-specific shortcuts, achieving within-cohort performance that collapses once the shortcut is removed. This is exactly the failure mode leave-one-farm-out is designed to expose, and exposing it is a principal output of the framework, because within-cohort validation cannot separate acoustic generalization from learned farm correlations. The confidence-stratified review makes the mechanism concrete: several of the highest-confidence errors for is_distress_related were maternal and social calls recorded in milking zones at Farms 1 and 3, where the model had learned to associate milking-zone context with distress patterns and confidently mislabelled an acoustically similar maternal contact call. This accords with the established view that animal vocalizations are context-conditioned signals whose interpretation cannot be decoupled from the conditions of production (Briefer, 2012; Manteuffel et al., 2004; Coutant et al., 2024).

### 7.4 Why reliability-aware interpretation is more informative than plain classification

Calibration and selective prediction are established techniques, but their systematic, target-specific integration in livestock acoustic modelling has not, to our knowledge, been explored before. Layer 3 showed that these are not optional post-processing steps but instruments that reveal structure plain labels conceal. For the non-vocal target, a leave-one-farm-out macro-F1 of 0.763 summarises discrimination, but calibration adds a separate dimension: expected calibration error fell from 0.087 to 0.023, so a predicted probability of 0.80 now sits much closer to the empirical frequency of correct predictions - which is what a tool that flags clips for human review actually needs (Guo et al., 2017; Silva Filho et al., 2023). The distress-related target shows the opposite face of the same property: calibration improved the Brier score (0.171 to 0.115) but lowered macro-F1 by shifting posterior weight toward the majority class - more honest probabilities at the cost of discrimination, the tension between accuracy and probability quality that benchmarks often conflate (Kull et al., 2019). Selective prediction adds a third dimension: withholding the least-confident 14% of proxy_functional_v1 predictions raised retained macro-F1 from 0.407 to 0.430, the difference between a system that can say it lacks evidence and one that labels every clip regardless (Geifman and El-Yaniv, 2017). An end-to-end log-mel CNN trained under identical protocols (Supplementary Table S7) outperformed the framework on raw accuracy for the easier targets (leave-one-farm-out macro-F1 0.864 and 0.733 for is_nonvocal and is_distress_related, versus 0.763 and 0.642) but did not rescue the harder targets and produced no calibrated probabilities, no abstention, and an expected calibration error larger at every target, ranging from roughly 1.8-fold (proxy_functional_v1) to 6.4-fold (is_nonvocal). The framework therefore prioritises interpretability and reliability over peak accuracy; benchmarking against larger architectures is left to future work.

### 7.5 Why proxy-state framing is scientifically appropriate

Proxy-state framing is the scientifically appropriate description of what acoustic monitoring can establish without synchronized physiological ground truth: not a welfare diagnosis, but a calibrated statement that a recording’s token sequence is consistent with some behavioral contexts and not others, under the conditions seen in training. What this buys, beyond flat classification, is a differentiated evidence picture. Non-vocal physiological units separate cleanly from behavioral vocalizations; within the behavioral space, reproductive estrus calls form a relatively well-defined category, while resource-seeking and distress-related calls remain context-entangled. This graduated characterization is exactly what Coutant et al. (2024) called for in their review of farm-animal bioacoustics, and a flat classifier that treats all sound events equivalently cannot produce it. Because the acoustic vocabulary is discovered without reference to any behavioral annotation, it remains reusable as feeding schedules, milking timestamps, or sensor streams become available - separating structure discovery from meaning assignment makes the representation more durable than end-to-end systems that couple the two.

### 7.6 Implications for livestock AI and welfare monitoring

The reliability-aware design has three practical implications that distinguish it from flat classification. It identifies when a prediction is reliable enough to act on and when it should be deferred to a human, through calibration and selective prediction; it identifies which acoustic signals generalize across farms, through the context-audited cross-farm protocol; and it provides an inspectable evidence representation - the model-behaviour dictionary - that can be combined with other sensing channels rather than treated as an opaque label predictor. These properties support cautious use as an evidence channel within a larger monitoring system rather than as a standalone welfare classifier. The framework is not deployment-ready: cross-farm macro-F1 of 0.407 for the functional proxy and 0.500 for the resource-related proxy are informative as evidence but below practical thresholds, and a three-farm corpus without synchronized physiological measurement cannot support clinical validation (Section 8). What it offers is a preliminary methodological foundation. Cross-farm generalizability is repeatedly identified as a central unsolved problem in bovine bioacoustics (Kate and Neethirajan, 2026; Jobarteh et al., 2025), and this study addresses it directly with a farm-held-out protocol rather than the single-site designs that predominate. The model-behaviour dictionary is the most directly reusable contribution: a structured profile of how each acoustic unit associates with proxy-state evidence under calibrated conditions, suitable as the acoustically grounded channel in the multimodal fusion architectures proposed for cattle welfare (Berckmans, 2017; Neculai-Valeanu et al., 2025) but rarely instantiated. In the longer term it could serve as the acoustic evidence module of a digital twin that accumulates confidence-weighted updates into a welfare trajectory rather than event-level labels (Neculai-Valeanu et al., 2025), with appropriate honesty about what acoustic evidence alone can establish.

## 8. Limitations and Future Work

### 8.1 Absence of synchronized welfare ground truth

The most fundamental limitation of this study is the absence of synchronized physiological or behavioral measurements that would allow the proxy-state targets to be validated against direct welfare evidence. No cortisol samples, heart rate recordings, rumination sensor logs, or independent ethological observations were collected in temporal alignment with the audio clips. The behavioral annotation categories used to define proxy targets represent expert human judgment about what was observable and audible at the time of recording - they do not independently confirm the animal’s arousal, affective valence, or welfare score. A clip labeled as a distress-related call reflects annotator agreement that the acoustic and contextual evidence was consistent with a distress event; it does not provide the kind of concurrent physiological evidence that would be needed to assert that the animal was in a welfare-compromised state at that moment.

This absence shapes every interpretive claim in the paper. The proxy-state targets, however well-motivated biologically, remain proxies: operationally defined functional groupings derived from annotation categories rather than validated welfare states. The distinction matters particularly for the distress-related and resource-related targets, where the boundaries between behavioral categories are inherently ambiguous without physiological corroboration. Confidence scores provided by annotators (Cohen’s κ = 0.884 at main-category level, κ = 0.699 at subcategory level; Kate and Neethirajan, 2026) confirm that the annotation scheme is internally consistent, but internal consistency is not the same as ground-truth validity. The framework explicitly acknowledges this gap through proxy-state framing throughout, and the calibration and abstention analyses in Layer 3 improve the trustworthiness of predicted probabilities - but they cannot substitute for biological validation.

### 8.2 Corpus scale and farm sample size

While modest in scale, this is one of the few multi-farm, real-barn dairy vocalization corpora evaluated under an explicit cross-farm protocol. The 569 clips from three commercial farms are substantial by current bovine-bioacoustics standards but small for robust generalization claims: each farm is a single holdout fold, so every cross-farm claim rests on one unseen farm at a time. Generalization to farms with different herd sizes, management, housing, feeding systems, or acoustic environments - which vary widely across commercial dairying - cannot be assumed from three farms in a single region of Atlantic Canada.

Per-farm imbalance compounds this: Farm 2 contributes 265 of 569 clips (46.6%), concentrated in the Feeding zone, while Farm 3 contributes 120 clips with the highest proportion of distress events (Supplementary Tables S2 and S3). The leave-one-farm-out benchmark therefore does not test generalization across equally representative farms, and the limited sample prevents strong conclusions about the performance distribution across the broader commercial population.

### 8.3 Farm-sensitive and collapsing proxy targets

Cross-farm generalizability is selective, not universal. The non-vocal target reached a LOFO macro-F1 of 0.763, while the resource-related target fell to chance (0.500, balanced accuracy 0.500), consistently across all three held-out farms. The root cause is structural rather than a matter of training-set size: the within-cohort context gains for the resource and multiclass targets did not survive farm holdout, the signature of a model exploiting farm-specific correlations between recording environment and behavioral category rather than learning acoustic generalizations - exactly what the leave-one-farm-out protocol is designed to expose. The positive cross-farm result for is_nonvocal should therefore not be extrapolated to the other targets. Farm management routines, pen layouts, and behavioral patterns create context entanglement that acoustic models partly absorb as shortcuts, a limitation that richer recording coverage and metadata - not post-hoc calibration or abstention - would be needed to address. The collapse of the resource-related and multiclass proxies is best read as a field-level finding about acoustic-only inference in livestock systems, made visible by the context-audited protocol.

### 8.4 Context-locked acoustic units and confounded features

The Layer 1 bias audit found six of the 48 units highly farm-locked (dominant-farm proportion ≥ 0.90); these cannot serve as farm-invariant evidence even if they capture real environment-specific sounds. Layer 2’s high-bias-filtered vocabulary excluded the eight units highly locked across any context variable, but moderately locked units (0.80–0.90) were retained, and the 0.90 cutoff was operationally pragmatic rather than theoretically derived. A more rigorous approach - causal-graph confound adjustment or invariant risk minimization - would require more farms and more balanced zone coverage than this corpus provides.

### 8.5 Metadata limitations: what was not recorded

The structured metadata available for each clip - farm, barn zone, microphone, microphone placement context, recording date, behavioral category, and subcategory - provide a useful but incomplete picture of the contextual conditions under which each vocalization was produced. Several important contextual variables were not recorded or are not available in aligned form.

No individual cow identifiers are present in the corpus. Repeated vocalizations by the same animal cannot be distinguished from vocalizations by different animals, which means that within-clip sequence structure may partly reflect individual vocal idiosyncrasies rather than state-level behavioral patterns. No feeding schedules, milking queue data, or activity sensor outputs were aligned with clip timestamps, which prevents behavioral state inference models from accessing the management-routine context that would be most informative for resource-related and handling-related vocalizations. No parity, lactation stage, or age information is available, meaning that the models operate over a mixed population of heifers and multiparous cows whose known differences in vocalization structure (Gavojdian et al., 2024) are not separable in the current analysis.

The subcategory distribution is also strongly right-tailed, with 13 of 39 subcategories containing five or fewer clips. While the proxy-state recoding in Layer 2 partially addressed this by grouping sparse subcategories into broader functional targets, the sparsest subcategories - including Injury_Response_Call (1 clip), Maternal_Response_Call (1 clip), Proximity_Maintenance_Call (1 clip), and Restraint_Protest_Call (1 clip) - provide no meaningful statistical basis for cross-farm evaluation and were absorbed into majority classes during proxy recoding. The resulting loss of fine-grained behavioral specificity is an unavoidable consequence of working with an ecologically realistic but imbalanced corpus.

### 8.6 Calibration improves probability quality, not biological validity

Calibration improved the probability quality of the framework’s predictions - for is_nonvocal, expected calibration error fell from 0.087 to 0.023 under the LOFO bundle - but it cannot convert a proxy-state prediction into a welfare assessment. The calibrators were fit and evaluated against the behavioral annotation labels, so improved calibration means predicted probabilities align more faithfully with annotation-label correctness, not that the model more reliably identifies welfare-relevant internal states. A model that is well-calibrated for the proxy targets can still be systematically wrong about underlying welfare if the annotation labels are themselves imprecise, as acknowledged throughout this study.

### 8.7 The model-behaviour dictionary is evidence-based, not semantically validated

The Layer 3 model-behaviour dictionary assigns reliability tiers to each of the 47 profiled acoustic units based on their within-corpus confidence distributions. The apparent paradox in these tiers - is_resource_related shows 95.7% of its tokens in the HIGH tier while achieving a LOFO macro-F1 of only 0.500 - is the clearest demonstration of the dictionary’s scope and limits. HIGH tier assignment reflects that the model assigns consistent, confident probabilities to clips containing those units within the training corpus. It does not reflect that those confidence assignments generalize across farms or that they carry validated biological meaning.

The dictionary’s token-level profiles should be read as structured summaries of model behavior under the current annotation scheme and corpus composition, not as biologically validated acoustic lexicon entries. Their practical value lies in making the model’s evidence structure inspectable - allowing analysts to identify which units dominate high-confidence correct predictions, which units are polysemous or context-sensitive, and which units are biologically plausible candidates for deeper investigation. They do not constitute ground truth for acoustic-behavioral mapping, and any future use of the dictionary for on-farm decision support would require independent biological validation against synchronized welfare measurements before claims about individual animal states could be justified.

### 8.8 Future work

The most direct path to strengthening the framework is enrichment of the contextual input. Aligning audio recordings with feeding schedule timestamps, milking queue logs, and activity sensor data from wearable collars would allow behavioral state models to use the same contextual variables that farm managers rely on for welfare assessment, replacing the indirect and confounded proxy of barn zone with direct behavioral state information. Synchronizing audio with cortisol, heart rate, or rumination sensor outputs for a subset of clips would allow at least some of the proxy-state targets to be validated against biological ground truth, transforming the current proxy-state framework into a proper supervised welfare model for those validated subsets.

Expanding the farm sample is equally important. Results from three farms can establish proof of concept for cross-farm acoustic generalization but cannot support strong claims about external validity. A future corpus spanning five or more farms across diverse management systems, housing designs, and geographic regions would allow the leave-one-farm-out evaluation to test true population-level generalization rather than single-farm holdout performance. The variability exposed in the current study - particularly for is_resource_related, where per-farm performance ranged from near-chance to moderate - suggests that more farms would reveal a richer picture of where acoustic generalization is stable and where it breaks down.

At the representation level, Layer 1 used a frozen encoder pretrained on human speech as a deliberate design choice rather than a default. The frozen encoder isolates the contribution of the downstream framework and evaluation protocol from confounding improvements that domain-specific fine-tuning would introduce. Future work may explore domain-specific fine-tuning, but the present study focuses on evaluating framework-level generalization rather than maximizing predictive performance. Fine-tuning the encoder on a larger bovine vocalization corpus, or applying animal-specific SSL models such as AVES (Hagiwara, 2023) or models pretrained directly on livestock sounds, could produce frame representations that capture bovine-specific acoustic structure more precisely than the current general-purpose backbone. This would not change the segmentation, clustering, and token-assignment logic but would provide richer raw material for acoustic unit discovery.

Individual cow tracking across recording sessions would address one of the current corpus’s most significant interpretive gaps. Identifying individual animals acoustically - via voice recognition techniques validated for cattle (Gavojdian et al., 2024) - would allow longitudinal vocal trajectories to be constructed, supporting welfare-relevant inference about changes in an individual’s vocal behavior over time rather than cross-sectional snapshots of population-level patterns. Finally, scaling the annotation process through semi-automated detection pipelines, with active learning-based selection of clips for expert review, would reduce the bottleneck that currently limits corpus expansion to the pace of manual segmentation and labeling.

## 9. Conclusions

Current approaches to livestock acoustic analysis typically treat the problem as a supervised classification task evaluated within a single recording environment. Such designs risk learning farm-specific correlations rather than generalizable acoustic structure and provide no explicit assessment of prediction reliability. This study addresses these limitations by introducing a reliability-aware computational framework that separates acoustic representation, proxy-state inference, and uncertainty evaluation, and by adopting a farm-held-out, context-audited evaluation protocol to expose shortcut learning in precision livestock systems.

Applied to 569 barn recordings from three commercial dairy farms, the framework discovered a 48-unit acoustic vocabulary without behavioral labels, tested four proxy-state targets under audio-only, context-conditioned, and leave-one-farm-out conditions, and assessed the reliability of the resulting predictions through calibration, selective prediction, and a model-behaviour dictionary.

The framework is not a deployment-ready welfare monitoring system. It does not have access to synchronized physiological ground truth, it was trained on three farms, and several of its proxy targets do not generalize reliably across recording sites. What it does establish is a methodological foundation: that self-supervised acoustic unit discovery can recover stable, biologically plausible structure from commercial dairy barn recordings without supervision, that proxy-state meaning can be attached cautiously to that structure with explicit cross-farm honesty checks, and that reliability-aware outputs - calibrated probabilities, abstention maps, and a confidence-stratified dictionary - are both feasible and informative in this domain. As recording corpora grow, synchronized metadata becomes available, and farm coverage expands, the layered architecture described here provides a principled path from acoustic structure discovery to validated welfare evidence modeling.

## Acknowledgements

The authors sincerely thank the Dairy Farmers of Nova Scotia and the Dairy Farmers of New Brunswick for generously providing access to their dairy farms and for their technical assistance and support throughout the data collection process. The authors also thank Jean Lynds (Operations Manager), Michael McConkey (Farm Manager), and Stewart Yuill (Animal Caretaker) for their support during data collection at the Ruminant Animal Centre, Dalhousie University.

## Ethics statement

All experimental procedures were reviewed and approved by the Dalhousie University Animal Ethics Committee (Protocol No. 2024-026). The study was conducted in accordance with local legislation and institutional requirements.

## Data availability

The curated audio dataset and associated metadata are deposited in Zenodo under restricted access (DOI: 10.5281/zenodo.17764250), with access provided through a Data Access Agreement to protect farm confidentiality.

