## Supplementary Material for "Beyond Accuracy: Reliability-Aware Cross-Farm Evaluation of Dairy Cow Vocalization Models"

**Supplementary Methods**

**S1. Segmentation refinement scorecard.** Candidate segmentations were ranked by a five-criterion weighted scorecard applied independently to each candidate: single-segment improvement, i.e. the reduction in the proportion of single-segment clips (weight 0.30); fragment control, i.e. avoidance of tiny sub-threshold fragments (0.20); downstream clustering quality under the K-study, measured by pairwise Adjusted Rand Index and the normalized selection score (0.20); sequence richness, measured by mean unique units per clip, normalized token entropy, and transition rate (0.20); and confound control, i.e. the amount of context locking introduced (0.10). Three penalties were applied: for failing to reduce single-segment clips, for fragment explosion (more than 25% of segments below 0.15 s, or a jump of more than seven percentage points from baseline), and for worsening context confounding by more than five percentage points.

**S2. K-study configuration and metrics.** For each candidate K, five independent MiniBatchKMeans runs were performed with seeds 7, 19, 31, 43, and 55 (batch size 256, maximum 300 iterations, 10 re-initialisations per run). Four internal metrics were averaged across seeds — silhouette score, Calinski–Harabasz score, Davies–Bouldin score, and cluster-balance statistics (minimum cluster size, median cluster size, normalized cluster entropy, singleton count) — and combined by normalized composite scoring with an explicit penalty for singleton or very small clusters. At K = 48 the mean silhouette score was 0.101, Calinski–Harabasz 166.85, Davies–Bouldin 2.196, mean minimum cluster size 13.4, normalized cluster entropy 0.964, and singleton count zero. The final clustering model was fit with K = 48 and fixed seed 31.

**S3. Calibration candidate sets and selection safeguards.** Calibration was selected independently for each target under four safeguards. (1) Candidates were target-appropriate: for the binary targets (is_nonvocal, is_distress_related, is_resource_related), raw probabilities, sigmoid/Platt scaling, isotonic regression, and temperature scaling; for the multiclass target (proxy_functional_v1), raw probabilities, multiclass temperature scaling, one-vs-rest Platt, and one-vs-rest isotonic. (2) Selection was made exclusively on the primary LOFO benchmark bundle, not on within-cohort random cross-validation, so that a method performing well only under random CV could not be chosen. (3) The selection criterion was probability quality — expected calibration error, Brier score, and log loss — combined by composite ranking; hard-label accuracy and macro-F1 were not selection criteria. (4) Calibration was fit group-aware: the calibrator applied to a held-out farm was fit only on the remaining farms' predictions.

**S4. Dictionary tier-threshold derivation.** Reliability tiers were assigned per token and per target from three criteria applied jointly: mean raw confidence across clips containing the token, the fraction of those clips classified as uncertain, and the number of clips contributing evidence. The tier boundaries (HIGH, MEDIUM, LOW, UNCERTAIN) were derived from the selective-prediction operating-point analysis and set target-specifically, because the four targets occupy different probability scales. The source was the validated token-sequence field of the Layer 2 master table (mean sequence length 8.37, maximum 12, 48 unique tokens in the corpus).

**Supplementary Tables**

**Supplementary Table S1. Complete subcategory-level clip distribution across all nine main behavioral categories and 39 subcategories (n = 569 clips).**

| **Main category** | **Subcategory** | **Clips** | **% of corpus** |
| --- | --- | --- | --- |
| Distress & Pain | General_Discomfort_Call | 11 | 1.9% |
| Distress & Pain | Pain_Related_Call | 11 | 1.9% |
| Distress & Pain | High_Frequency_Distress | 10 | 1.8% |
| Distress & Pain | Illness_Indication_Call | 7 | 1.2% |
| Distress & Pain | Injury_Response_Call | 1 | 0.2% |
| Environmental & Situational | Novel_Environment_Call | 2 | 0.4% |
| Estrus & Mating Behavior | Estrus_Call | 117 | 20.6% |
| Estrus & Mating Behavior | Mating_Excitement_Call | 10 | 1.8% |
| Feeding & Hunger Related | Feed_Anticipation_Call | 112 | 19.7% |
| Feeding & Hunger Related | Post_Feeding_Satisfaction | 9 | 1.6% |
| Feeding & Hunger Related | Empty_Feeder_Call | 6 | 1.1% |
| Feeding & Hunger Related | Hunger_Frustration_Call | 6 | 1.1% |
| Feeding & Hunger Related | Feed_Quality_Response | 5 | 0.9% |
| Feeding & Hunger Related | Feed_Competition_Call | 3 | 0.5% |
| Maternal & Calf Communication | Mother_Separation_Call | 9 | 1.6% |
| Maternal & Calf Communication | Calf_Contact_Call | 6 | 1.1% |
| Maternal & Calf Communication | Maternal_Response_Call | 1 | 0.2% |
| Milking & Handling | Pre_Milking_Call | 12 | 2.1% |
| Milking & Handling | Milking_Discomfort_Call | 9 | 1.6% |
| Milking & Handling | Handling_Stress_Call | 3 | 0.5% |
| Milking & Handling | Restraint_Protest_Call | 1 | 0.2% |
| Non-Vocal Sounds | Breathing_Respiratory_Sounds | 77 | 13.5% |
| Non-Vocal Sounds | Chewing_Rumination_Sounds | 31 | 5.4% |
| Non-Vocal Sounds | Drinking_Slurping_Sounds | 17 | 3.0% |
| Non-Vocal Sounds | Cough_Calls | 11 | 1.9% |
| Non-Vocal Sounds | Licking_Sounds | 3 | 0.5% |
| Non-Vocal Sounds | Movement_Associated_Sounds | 3 | 0.5% |
| Social Recognition & Interaction | Greeting_Call | 11 | 1.9% |
| Social Recognition & Interaction | Social_Bonding_Call | 9 | 1.6% |
| Social Recognition & Interaction | Herd_Coordination_Call | 7 | 1.2% |
| Social Recognition & Interaction | Individual_Recognition_Call | 7 | 1.2% |
| Social Recognition & Interaction | Response_Exchange_Call | 7 | 1.2% |
| Social Recognition & Interaction | Group_Contact_Call | 6 | 1.1% |
| Social Recognition & Interaction | Proximity_Maintenance_Call | 1 | 0.2% |
| Water & Thirst Related | Thirst_Anticipation_Call | 9 | 1.6% |
| Water & Thirst Related | Water_Frustration_Call | 8 | 1.4% |
| Water & Thirst Related | Dehydration_Distress_Call | 5 | 0.9% |
| Water & Thirst Related | Water_Quality_Response | 4 | 0.7% |
| Water & Thirst Related | Drinking_Competition_Call | 2 | 0.4% |
| **Total** |  | **569** | **100%** |

**Supplementary Table S2. Cross-tabulation of clip counts by farm and main behavioral category for farm-identified clips (n = 559).**

| **Farm** | **Distress & Pain** | **Envir. & Sit.** | **Estrus & Mating** | **Feeding & Hunger** | **Maternal & Calf** | **Milking & Handling** | **Non-Vocal** | **Social Recognition** | **Water & Thirst** | **Total** |
| --- | --- | --- | --- | --- | --- | --- | --- | --- | --- | --- |
| Farm 1 | 10 | 2 | 61 | 10 | 3 | 12 | 38 | 29 | 9 | 174 |
| Farm 2 | 4 | 0 | 25 | 129 | 0 | 1 | 81 | 9 | 16 | 265 |
| Farm 3 | 26 | 0 | 41 | 2 | 10 | 12 | 22 | 4 | 3 | 120 |
| **Total** | **40** | **2** | **127** | **141** | **13** | **25** | **141** | **42** | **28** | **559** |

**Supplementary Table S3. Cross-tabulation of clip counts by barn zone and main behavioral category for farm-identified clips (n = 559).**

| **Barn zone** | **Distress & Pain** | **Envir. & Sit.** | **Estrus & Mating** | **Feeding & Hunger** | **Maternal & Calf** | **Milking & Handling** | **Non-Vocal** | **Social Recognition** | **Water & Thirst** | **Total** |
| --- | --- | --- | --- | --- | --- | --- | --- | --- | --- | --- |
| Feeding | 18 | 0 | 35 | 133 | 0 | 0 | 68 | 14 | 0 | 268 |
| Resting | 9 | 1 | 61 | 8 | 3 | 0 | 52 | 10 | 0 | 144 |
| Drinking | 7 | 0 | 31 | 0 | 0 | 0 | 8 | 4 | 12 | 62 |
| Milking | 6 | 1 | 0 | 0 | 10 | 25 | 0 | 12 | 0 | 54 |
| Water Station | 0 | 0 | 0 | 0 | 0 | 0 | 13 | 2 | 16 | 31 |
| **Total** | **40** | **2** | **127** | **141** | **13** | **25** | **141** | **42** | **28** | **559** |

**Supplementary Table S4. Clip duration summary statistics (mean, median, minimum, and maximum in seconds) by main behavioral category (n = 569 clips).**

| **Main category** | **Clips** | **Mean (s)** | **Median (s)** | **Min (s)** | **Max (s)** |
| --- | --- | --- | --- | --- | --- |
| Non-Vocal Sounds | 142 | 24.5 | 18.7 | 2.8 | 444.9 |
| Feeding & Hunger Related | 141 | 16.5 | 13.4 | 9.9 | 56.7 |
| Estrus & Mating Behavior | 127 | 12.9 | 13.4 | 3.6 | 15.0 |
| Social Recognition & Interaction | 48 | 36.6 | 15.8 | 10.1 | 267.0 |
| Distress & Pain | 40 | 17.4 | 15.2 | 10.3 | 38.2 |
| Water & Thirst Related | 28 | 22.1 | 19.2 | 10.1 | 57.8 |
| Milking & Handling | 25 | 24.0 | 23.4 | 20.2 | 35.4 |
| Maternal & Calf Communication | 16 | 49.6 | 16.0 | 11.1 | 321.0 |
| Environmental & Situational | 2 | 32.9 | 32.9 | 32.2 | 33.7 |
| **All clips** | **569** | **21.0** | **13.8** | **2.8** | **444.9** |

**Supplementary Table S5. Summary metrics for the official refined Layer 1 acoustic unit discovery pipeline, including segmentation, clustering, and dictionary scaffold statistics.**

| **Metric** | **Value** |
| --- | --- |
| n_clips_ssl | 569 |
| n_clips_segmented | 569 |
| n_segments_total | 4764 |
| mean_clip_duration_sec | 21.038 |
| median_clip_duration_sec | 13.808 |
| mean_segments_per_clip | 8.373 |
| median_segments_per_clip | 9 |
| single_segment_clips | 6 |
| single_segment_clip_pct | 1.054 |
| max_segments_single_clip | 12 |
| n_units | 48 |
| prototypes_total | 144 |
| prototypes_per_unit | 3 |

**Supplementary Table S6. Context-bias audit summary for all 48 discovered acoustic units across four metadata fields (farm, barn zone, microphone, and microphone placement context), reporting flagged and high-bias unit counts and dominant context proportions.**

| **Field Name** | **Flagged Units** | **High Bias Units** | **Median Dominant Context Proportion** | **Max Dominant Context Proportion** |
| --- | --- | --- | --- | --- |
| farm | 9 | 6 | 0.611128049 | 1 |
| barn_zone | 10 | 4 | 0.548809524 | 1 |
| microphone | 9 | 2 | 0.510706914 | 1 |
| mic_placement_context | 4 | 1 | 0.487625139 | 1 |

**Supplementary Table S7. End-to-end CNN baseline comparison across the four proxy-state targets under matched 5-fold cross-validation (audio-only) and leave-one-farm-out evaluation. CNN ECE values are raw, as the baseline applies no post-hoc calibration.**

| **Target** | **Framework 5-fold macro-F1** | **CNN 5-fold macro-F1 (mean ± std)** | **Framework LOFO macro-F1** | **CNN LOFO macro-F1 (mean ± std)** | **Framework ECE (LOFO)** | **CNN raw ECE (LOFO, mean)** |
| --- | --- | --- | --- | --- | --- | --- |
| is_nonvocal | 0.808 | 0.921 ± 0.039 | 0.763 | 0.864 ± 0.018 | 0.023 | 0.148 |
| is_distress_related | 0.689 | 0.797 ± 0.032 | 0.642 | 0.733 ± 0.092 | 0.116 | 0.229 |
| is_resource_related | 0.704 | 0.664 ± 0.050 | 0.500 | 0.439 ± 0.051 | 0.073 | 0.199 |
| proxy_functional_v1 | 0.654 | 0.549 ± 0.041 | 0.407 | 0.416 ± 0.142 | 0.099 | 0.175 |

**Supplementary Table S8. Segmentation candidate parameter configurations. Key outcomes reported for the selected candidate (Candidate C).**

| **Parameter** | **Baseline** | **Candidate A (gentle)** | **Candidate B (balanced)** | **Candidate C (richer)** |
| --- | --- | --- | --- | --- |
| Smoothing window (frames) | 5 | 5 | 5 | 4 |
| Change-score smoothing (frames) | 3 | 3 | 2 | 2 |
| Peak prominence | 0.025 | 0.022 | 0.020 | 0.018 |
| MAD multiplier | 3.0 | 2.75 | 2.50 | 2.25 |
| Min inter-peak distance (s) | 0.20 | 0.18 | 0.15 | 0.14 |
| Min segment duration (s) | 0.25 | 0.20 | 0.18 | 0.16 |
| Max segments per clip | None | None | None | 12 |
| Single-segment clips (%) | 81.7% | - | - | 1.1% |
| Total segments | 819 | - | - | 4,764 |
| Tiny fragments < 0.15s (%) | - | - | - | 0% |

*Note: Intermediate outcome statistics for Candidates A and B were computed during the refinement sweep but are not reported here because selection was made by composite scorecard ranking. Only the selected candidate (Candidate C) and the baseline are shown in full.*

**Supplementary Table S9. K-study summary results for the five tested K values (means across five seeds).**

| **K** | **Mean pairwise ARI** | **Mean silhouette** | **Mean CH score** | **Mean DB score** | **Mean min cluster size** | **Normalized entropy** | **Mean singletons** |
| --- | --- | --- | --- | --- | --- | --- | --- |
| 32 | 0.571 | 0.112 | 189.43 | 2.012 | 28.6 | 0.945 | 0.0 |
| 48 | 0.501 | 0.101 | 166.85 | 2.196 | 13.4 | 0.964 | 0.0 |
| 64 | 0.443 | 0.093 | 148.21 | 2.387 | 7.8 | 0.971 | 0.4 |
| 80 | 0.391 | 0.088 | 131.55 | 2.541 | 4.2 | 0.976 | 1.2 |
| 96 | 0.341 | 0.082 | 119.30 | 2.703 | 2.8 | 0.979 | 3.4 |

**Supplementary Table S10. Context bias audit summary: flagged and highly locked units per audited field.**

| **Context variable** | **Flagged units (≥ 0.80)** | **Highly locked units (≥ 0.90)** | **Median dominant proportion** | **Maximum dominant proportion** |
| --- | --- | --- | --- | --- |
| Farm | 9 | 6 | 0.611 | 1.000 |
| Barn zone | 10 | 4 | 0.549 | 1.000 |
| Microphone | 9 | 2 | 0.511 | 1.000 |
| Mic placement context | 4 | 1 | 0.488 | 1.000 |

**Supplementary Table S11. Summary of Layer 1 outputs: acoustic vocabulary, segmentation statistics, and bias audit indicators.**

| **Output** | **Value** |
| --- | --- |
| Clips in corpus | 569 |
| SSL embedding dimension | 768 |
| Frame hop | ~0.020s |
| Total segments (refined) | 4,764 |
| Mean segments per clip | 8.37 |
| Median segments per clip | 9 |
| Maximum segments per clip | 12 |
| Unit inventory size (K) | 48 |
| Total token events | 4,764 |
| Top 15 units share | 54% of all events |
| Prototype segments per unit | 3 |
| Mean pairwise ARI at K=48 | 0.501 |
| Silhouette at K=48 | 0.101 |
| Singleton clusters | 0 |
| Farm-level highly locked units | 6 |
| Farm-level flagged units | 9 |
| Barn-zone flagged units | 10 |

**Supplementary Table S12. Best audio-only results by proxy-state target (class-balanced logistic regression, 5-fold stratified cross-validation, 569 clips).**

| **Target** | **Best feature family** | **Macro-F1** | **Balanced accuracy** |
| --- | --- | --- | --- |
| is_nonvocal | TF-IDF 1–3-gram (bias-filtered) | 0.808 | 0.846 |
| proxy_functional_v1 | TF-IDF 1–3-gram (full) | 0.654 | 0.674 |
| is_resource_related | Unigram composition (full) | 0.704 | 0.725 |
| is_distress_related | TF-IDF 1–3-gram (full) | 0.689 | 0.736 |

**Supplementary Table S13. Best audio-plus-context results and gains over audio-only (5-fold cross-validation, 569 clips, full context).**

| **Target** | **Best audio-only Macro-F1** | **Best audio + context Macro-F1** | **Δ Macro-F1** | **Best audio + context Bal. Acc** |
| --- | --- | --- | --- | --- |
| is_nonvocal | 0.808 | 0.847 | +0.039 | 0.882 |
| proxy_functional_v1 | 0.654 | 0.766 | +0.112 | 0.772 |
| is_distress_related | 0.689 | 0.800 | +0.111 | 0.866 |
| is_resource_related | 0.704 | 0.859 | +0.155 | 0.882 |

**Supplementary Table S14. Per-farm holdout spread statistics for pooled LOFO macro-F1 by proxy-state target.**

| **Target** | **Best LOFO bundle** | **Mean F1 (per farm)** | **Std** | **Range (min–max)** |
| --- | --- | --- | --- | --- |
| is_nonvocal | TF-IDF 1–3-gram (filtered), audio-only | 0.735 | 0.046 | 0.674–0.785 |
| proxy_functional_v1 | Transition probs (full), recording-env. context | 0.412 | 0.093 | 0.286–0.510 |
| is_distress_related | Transition probs (filtered), recording-env. context | 0.598 | 0.014 | 0.579–0.610 |
| is_resource_related | Unigram composition (full), audio-only | 0.459 | 0.010 | 0.445–0.468 |

**Supplementary Table S15. Qualitative review summary: accuracy and confidence statistics by target (LOFO best bundle per target, 559 known-farm clips).**

| **Target** | **Accuracy** | **Mean confidence** | **Confidence: correct** | **Confidence: wrong** |
| --- | --- | --- | --- | --- |
| is_nonvocal | 0.809 | 0.393 | 0.352 | 0.566 |
| is_distress_related | 0.807 | 0.316 | 0.262 | 0.542 |
| is_resource_related | 0.578 | 0.430 | 0.389 | 0.487 |
| proxy_functional_v1 | 0.442 | 0.535 | 0.654 | 0.442 |

**Supplementary Table S16. Model-behaviour dictionary reliability tier distribution by target (47 tokens evaluated per target).**

| **Target** | **HIGH (%)** | **MEDIUM (%)** | **LOW (%)** | **UNCERTAIN (%)** | **Polysemous candidates** | **Mean token confidence** | **Mean uncertain rate** |
| --- | --- | --- | --- | --- | --- | --- | --- |
| is_nonvocal | 29 (61.7%) | 12 (25.5%) | 4 (8.5%) | 2 (4.3%) | 5 | 0.693 | 0.232 |
| proxy_functional_v1 | 42 (89.4%) | 1 (2.1%) | 4 (8.5%) | 0 | 7 | 0.527 | 0.130 |
| is_distress_related | 43 (91.5%) | 4 (8.5%) | 0 | 0 | 2 | 0.789 | 0.104 |
| is_resource_related | 45 (95.7%) | 2 (4.3%) | 0 | 0 | 4 | 0.626 | 0.070 |

**Supplementary Table S17. Confidence-stratified review summary: accuracy, confidence, and abstention rate by target (primary LOFO benchmark, 559 known-farm clips per target).**

| **Target** | **Accuracy** | **Mean raw confidence** | **Mean calibrated confidence** | **Abstention rate** | **Mean entropy** |
| --- | --- | --- | --- | --- | --- |
| is_nonvocal | 0.809 | 0.722 | 0.808 | 0.182 | 0.560 |
| is_distress_related | 0.807 | 0.787 | 0.851 | 0.102 | 0.437 |
| is_resource_related | 0.578 | 0.633 | 0.633 | 0.079 | 0.643 |
| proxy_functional_v1 | 0.442 | 0.535 | 0.414 | 0.132 | 1.129 |

**Supplementary Table S18. Final per-target Layer 3 reliability summary.**

| **Target** | **Role** | **Calibration** | **LOFO macro-F1 (raw)** | **LOFO macro-F1 (calibrated)** | **Selective gain (80% cov.)** | **Dictionary HIGH tier (%)** | **Interpretation** |
| --- | --- | --- | --- | --- | --- | --- | --- |
| is_nonvocal | Flagship | Sigmoid/Platt | 0.763 | 0.717 | +0.030 | 61.7% | Strongest probability quality; raw abstention informative |
| proxy_functional_v1 | Main multiclass | Temperature | 0.407 | 0.407 | +0.023 | 89.4% | Calibration and abstention both yield modest gains |
| is_distress_related | Meaningful but mixed | Sigmoid/Platt | 0.642 | 0.483 | 0.0% | 91.5% | Probability quality improved; abstention not beneficial |
| is_resource_related | Stress test | Raw (kept) | 0.500 | 0.500 | +0.003 | 95.7% | No calibration help; high-tier dictionary entries within-corpus only |

**Supplementary Figures**

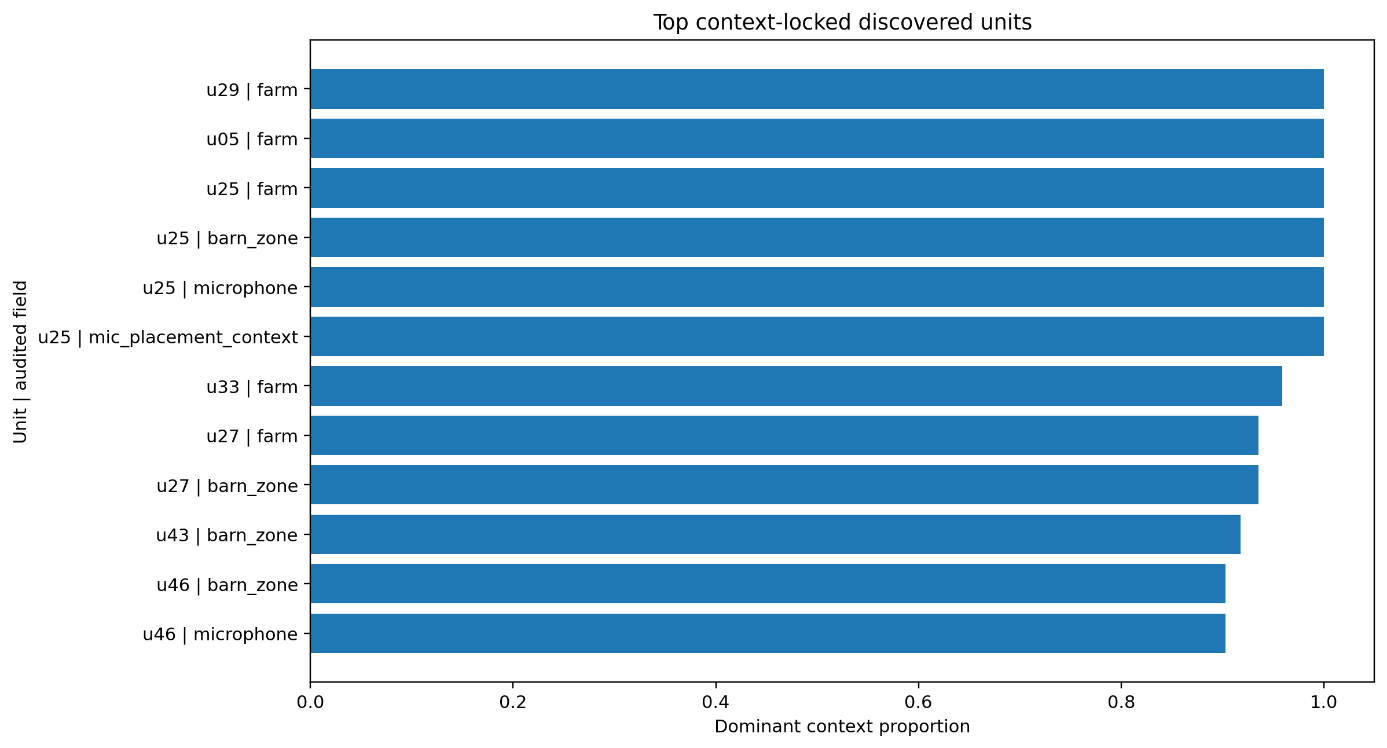

***Supplementary Figure S1. Top context-locked acoustic units with dominant context proportions****. Each bar represents a discovered unit flagged during the bias audit, with the proportion of segments concentrated in a single dominant context value. Units with high dominant proportions indicate potential confounding with recording conditions or farm-specific routines rather than generalizable acoustic structure.*

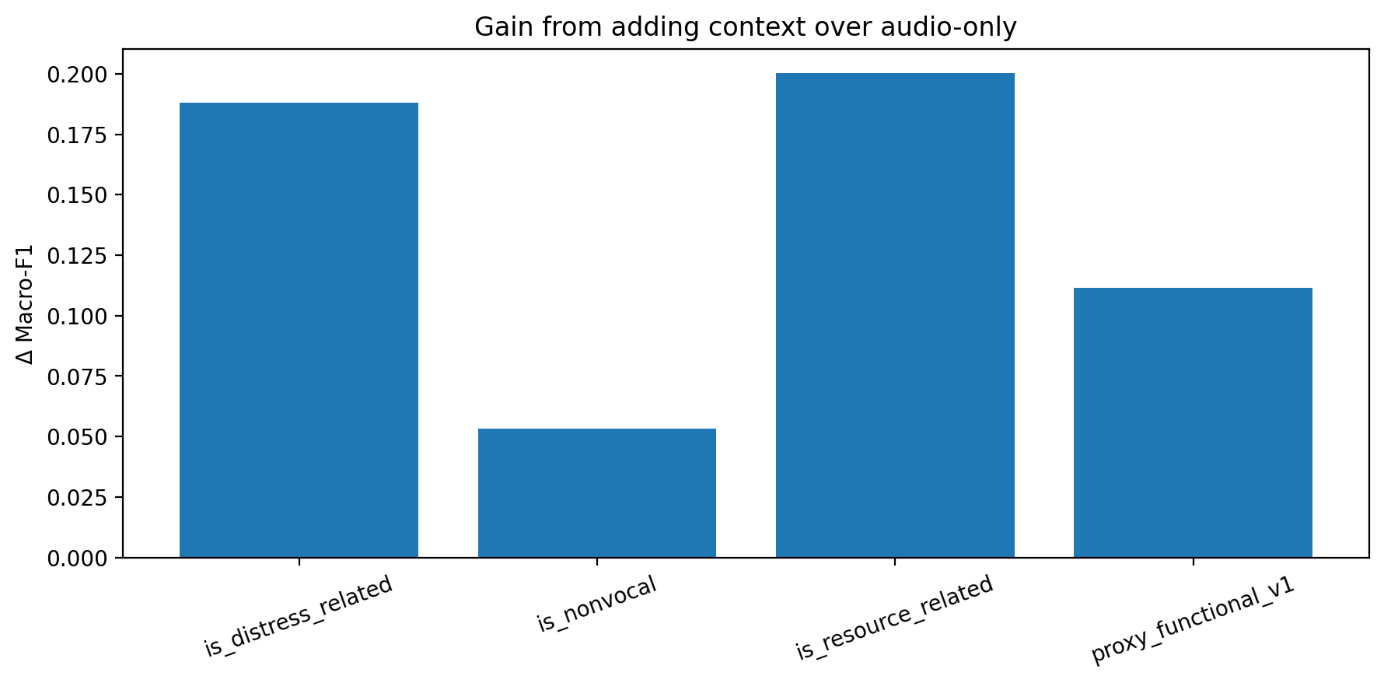

***Supplementary Figure S2. Per-target change in macro-F1 from the audio-only baseline to the audio-plus-context condition under matched cross-validation.*** *Positive values indicate that the inclusion of contextual metadata (farm, barn zone, microphone, microphone placement, day, and date) improved proxy-state inference. Targets with greater ambiguity (e.g., is_distress_related, is_resource_related) showed the largest context gains, while the acoustically self-contained is_nonvocal target improved only modestly.*

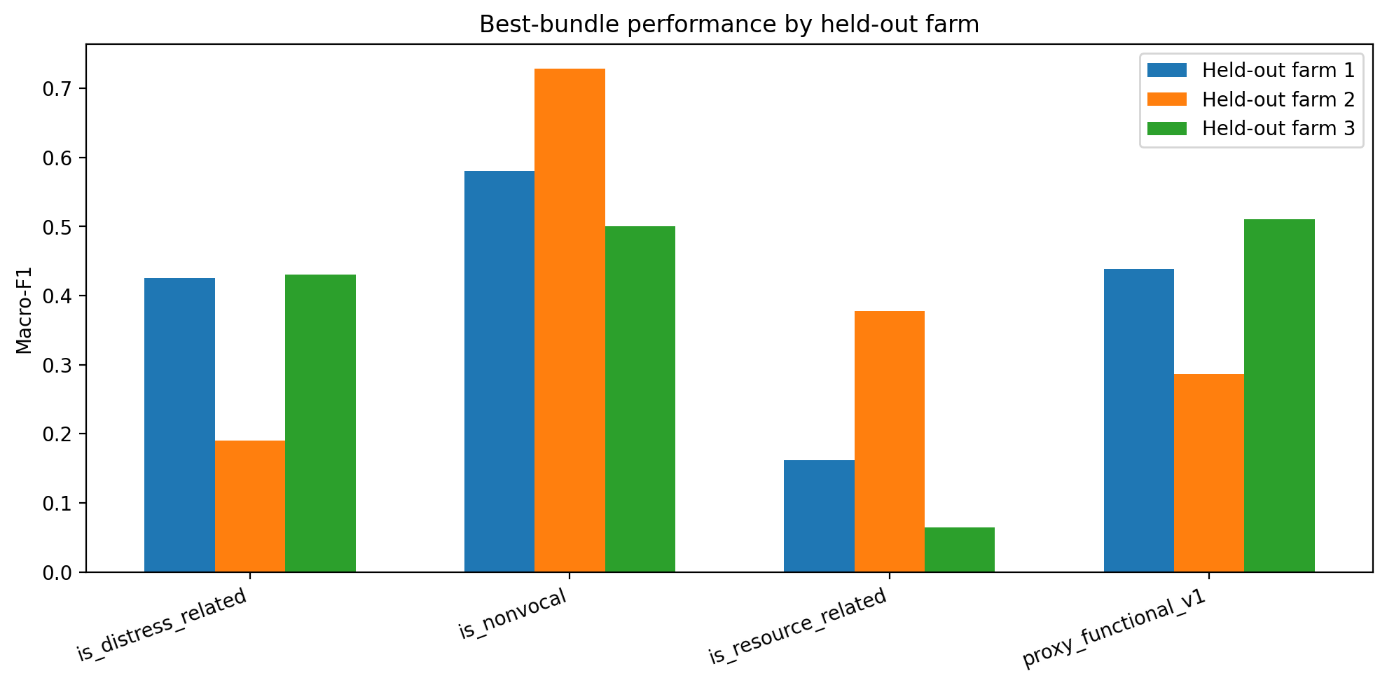

***Supplementary Figure S3. Per-farm holdout macro-F1 spread across proxy-state targets under leave-one-farm-out (LOFO) evaluation.*** *Each point represents the macro-F1 obtained when a single farm was held out entirely, and the range illustrates cross-farm variability. Targets with narrow spread indicate more farm-robust proxy-state signal; wider spread indicates greater farm sensitivity.*

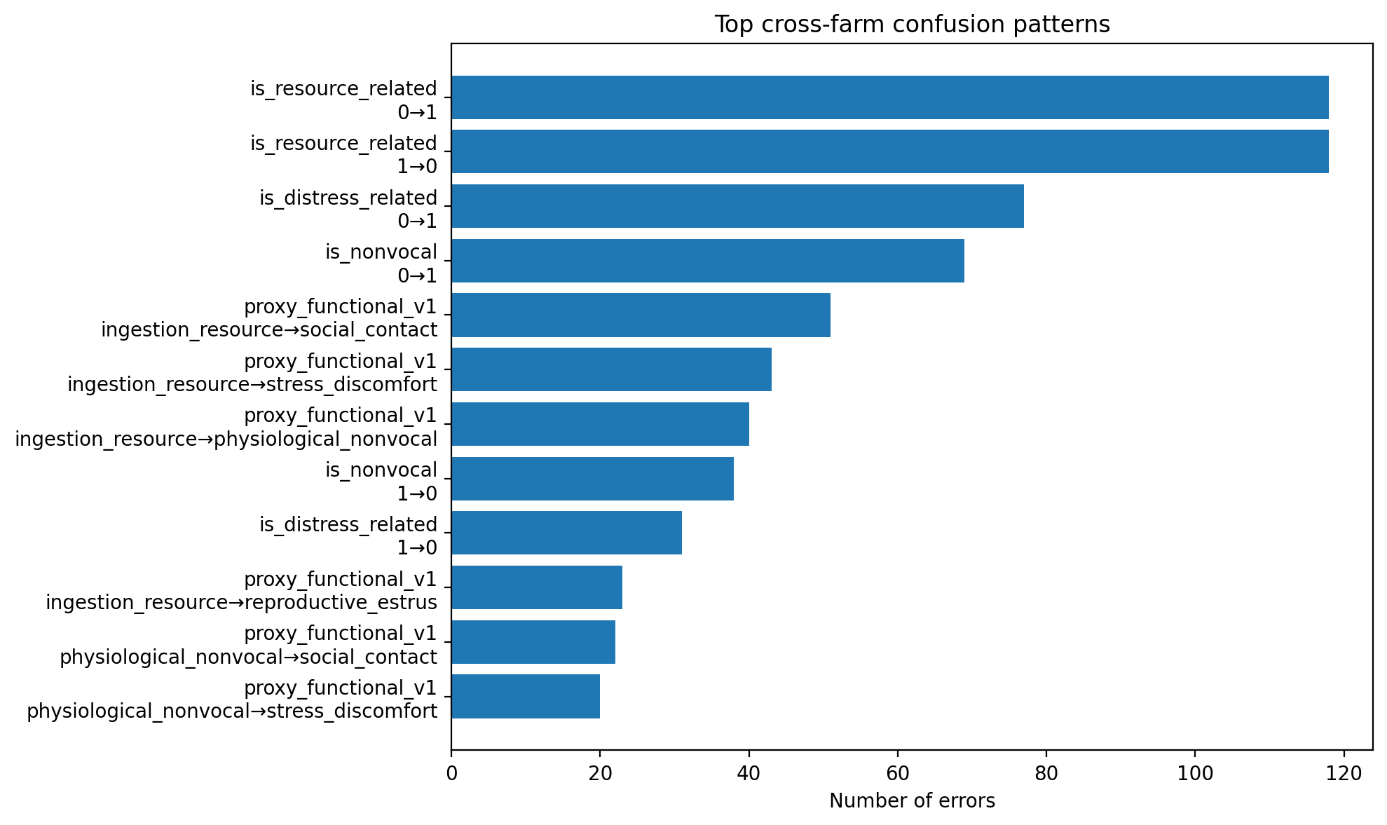

***Supplementary Figure S4. Top confusion patterns for the five-class proxy_functional_v1 target, showing the most frequent true-to-predicted label pairs and the dominant farm associated with each error type.*** *This figure highlights systematic misclassification tendencies, particularly between acoustically or contextually overlapping proxy-state classes such as ingestion/resource-related and social/contact categories.*

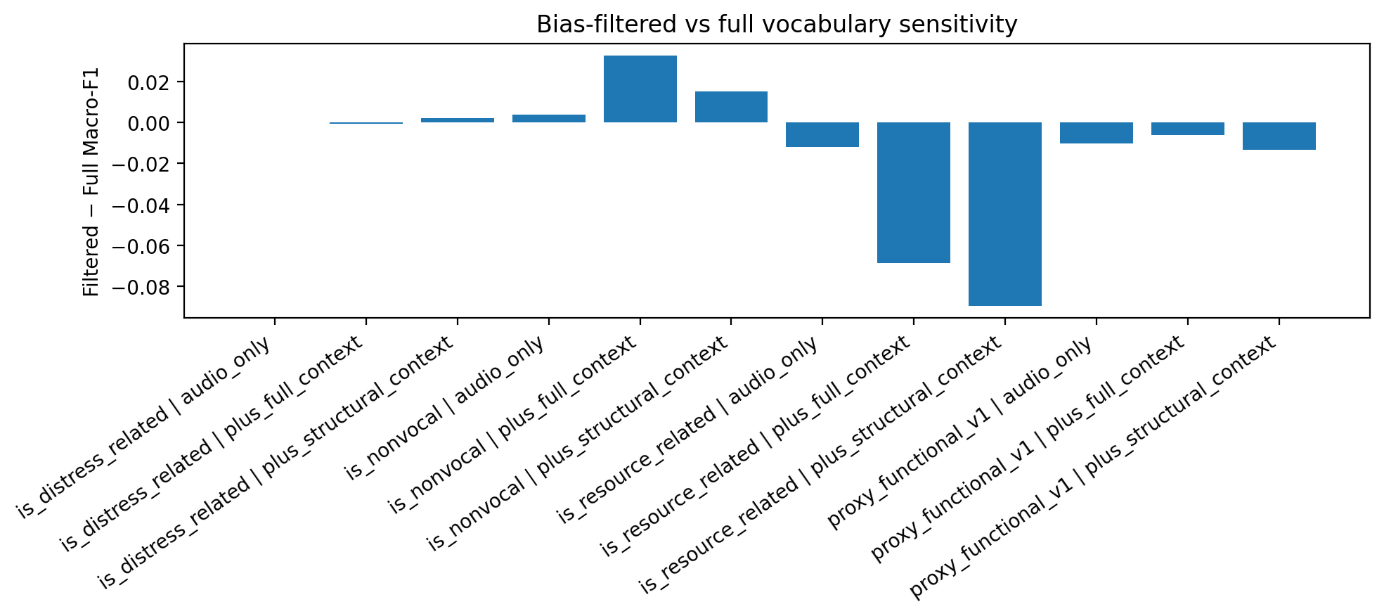

***Supplementary Figure S5. Bias-filtered versus full-vocabulary sensitivity under leave-one-farm-out cross-farm evaluation.*** *Each bar shows the macro-F1 difference (filtered minus full vocabulary) for each proxy-state target and context variant combination. Positive values indicate that removing high-bias-flagged acoustic units improved cross-farm generalization; negative values indicate that the full vocabulary performed better. The is_nonvocal target showed modest gains from filtering under context-enriched conditions, whereas is_resource_related under structural context showed the largest performance loss.*

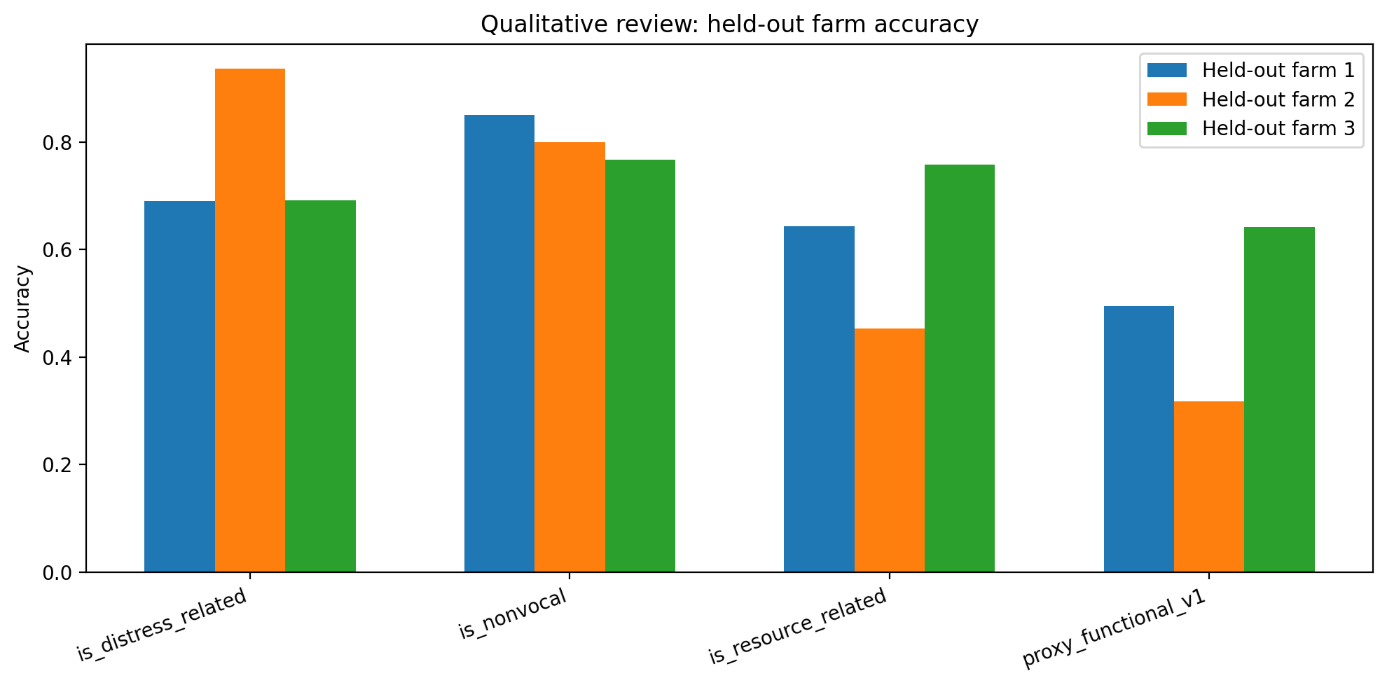

***Supplementary Figure S6. Per-farm held-out accuracy across proxy-state targets under leave-one-farm-out evaluation.*** *Grouped bars show accuracy when each of the three farms was held out entirely. The is_nonvocal target showed the most stable accuracy across held-out farms, while is_resource_related and proxy_functional_v1 showed substantial farm-dependent variability, with Farm 2 holdout producing the lowest accuracy for both targets.*

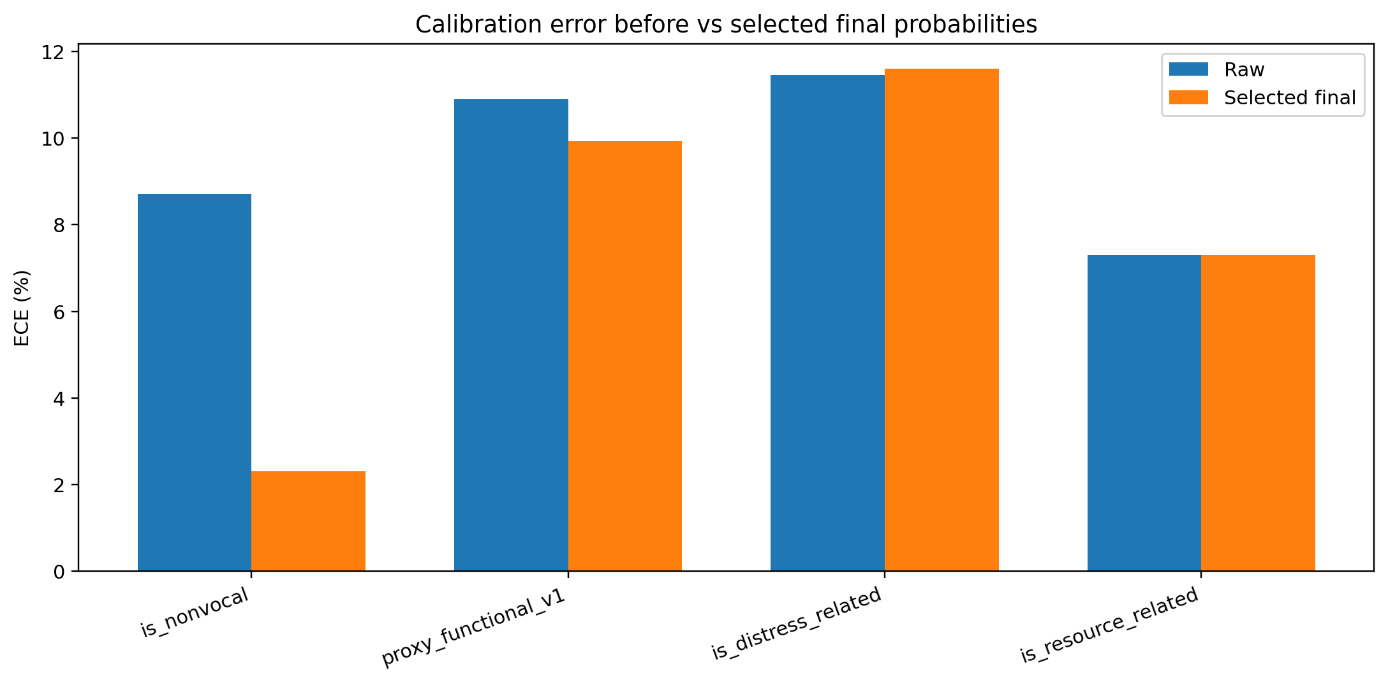

***Supplementary Figure S7. Calibration improvement summary showing raw versus selected expected calibration error (ECE) and Brier score for each proxy-state target under the primary LOFO benchmark bundle.*** *For each target, the selected calibration method (sigmoid/Platt scaling for is_nonvocal and is_distress_related, temperature scaling for proxy_functional_v1, raw preservation for is_resource_related) was chosen based on probability-quality criteria. Reductions in ECE and Brier score indicate improved alignment between predicted probabilities and observed outcomes.*

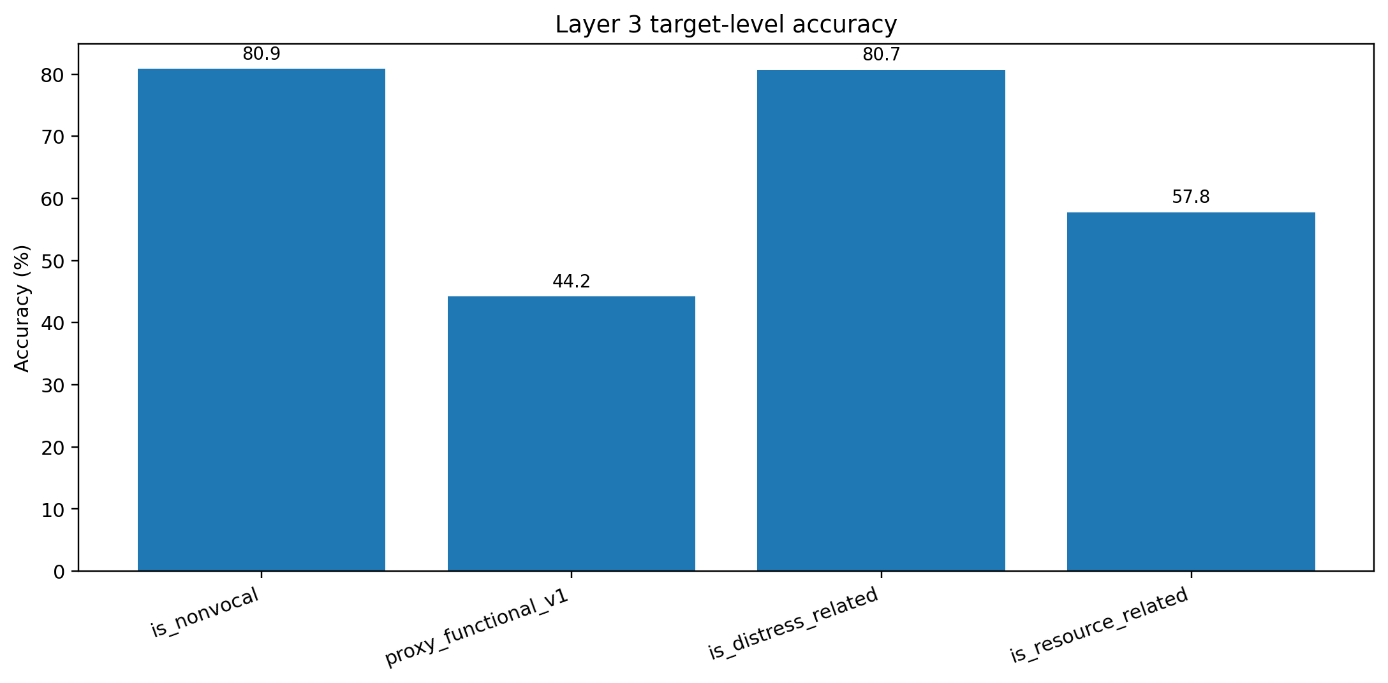

***Supplementary Figure S8. Per-target accuracy summary with abstention rate overlay under the selected operating point. For each proxy-state target, the left axis shows accuracy on the retained (non-abstained) subset, and the overlay indicates the proportion of clips deferred at the chosen confidence threshold.*** *Targets where abstention meaningfully improved retained-set quality (e.g., proxy_functional_v1) are distinguished from targets where abstention provided limited or no improvement (e.g., is_distress_related, is_resource_related).*
